# *ATP13A3* Variants Promote Pulmonary Arterial Hypertension by Disrupting Polyamine Transport

**DOI:** 10.1101/2023.08.29.554603

**Authors:** Bin Liu, Mujahid Azfar, Ekaterina Legchenko, James A. West, Shaun Martin, Chris Van den Haute, Veerle Baekelandt, John Wharton, Luke Howard, Martin R. Wilkins, Peter Vangheluwe, Nicholas W. Morrell, Paul D. Upton

**Affiliations:** Section of Cardio and Respiratory Medicine, Department of Medicine, Cambridge, UK; Department of Cellular and Molecular Medicine, KU Leuven, Leuven, Belgium; Cambridge Institute of Therapeutic Immunology and Infectious Disease, Jeffrey Cheah Biomedical Centre, Cambridge, UK; Division of Gastroenterology and Hepatology, Department of Medicine, Cambridge, UK; Department of Biochemistry and Cambridge Systems Biology Centre, University of Cambridge, Cambridge, UK; Laboratory for Neurobiology and Gene Therapy, Department of Neurosciences, KU Leuven, Leuven, Belgium; Leuven Viral Vector Core, KU Leuven, Leuven, Belgium; Department of Medicine, Imperial College London, London, UK

**Author notes:** joint first author (Bin Liu, Mujahid Azfar, Ekaterina Legchenko). joint senior authors (Peter Vangheluwe, Nicholas W. Morrell, Paul D. Upton). **Corresponding author:** Dr Paul D Upton PhD, Department of Medicine, Heart and Lung Research Institute, Papworth Road Cambridge CB2 0BB, UK. Bin Liu: Margaret Turner Warwick Centre for Fibrosing Lung Disease, National Heart and Lung Institute, Imperial College London, London, UK. Ekaterina Legchenko: Department of Cardiovascular Physiology, Heidelberg University, Medical Faculty Mannheim, Mannheim, Germany. Shaun Martin: Galapagos NV, Generaal De Wittelaan L11, 2800 Mechelen, Belgium.

## Abstract

**Aims:** Potential loss-of-function variants of *ATP13A3*, the gene encoding a P5B-type transport ATPase of undefined function, were recently identified in pulmonary arterial hypertension (PAH) patients. ATP13A3 is implicated in polyamine transport but its function has not been fully elucidated. Here, we sought to determine the biological function of ATP13A3 in vascular endothelial cells and how PAH-associated mutations may contribute to disease pathogenesis. We also generated mice harbouring an *Atp13a3* variant analogous to a human disease-associated variant to establish whether these mice develop PAH.

**Methods and Results:** We studied the impact of ATP13A3 deficiency and overexpression in endothelial cell (EC) models (human pulmonary ECs, blood outgrowth ECs (BOECs) and HMEC-1 cells), including a PAH patient-derived BOEC line harbouring an ATP13A3 variant (LK726X). ATP13A3 localised to the recycling endosomes of human ECs. Knockdown of ATP13A3 in ECs generally reduced the basal polyamine content, consistently reduced putrescine uptake, and altered the expression of enzymes involved in polyamine metabolism. Conversely, overexpression of wild-type ATP13A3 increased polyamine uptake, with an overall preference of putrescine > spermidine > spermine. Functionally, loss of ATP13A3 was associated with reduced EC proliferation, increased apoptosis in serum starvation and increased monolayer permeability to thrombin. Assessment of five PAH-associated missense ATP13A3 variants (L675V, M850I, V855M, R858H, L956P) confirmed loss-of-function phenotypes represented by impaired polyamine transport and dysregulated EC function. Furthermore, mice carrying a heterozygous germ-line *Atp13a3* frameshift variant representing a human mutation spontaneously developed a PAH phenotype, with increased pulmonary pressures, right ventricular remodelling and muscularisation of pulmonary vessels.

**Conclusion:** We identify ATP13A3 as a polyamine transporter, deficiency of which leads to EC dysfunction and predisposes to PAH. This suggests a need for targeted therapies to alleviate the imbalances in polyamine homeostasis and EC dysfunction in PAH.

**Translational perspective:** Rare missense *ATP13A3* disease-associated variants have been identified in patients with pulmonary arterial hypertension (PAH), though their pathogenicity has not been confirmed as the function of ATP13A3 is not known. We have identified ATP13A3 as a polyamine transporter, showing that ATP13A3 deficiency impaired polyamine homeostasis and uptake, and drove endothelial dysfunction. Conversely, overexpression increased polyamine uptake and rescued the proapoptotic phenotype of cells harbouring a disease-associate variant. Mice heterozygous for a disease-associated Atp13a3 mutation spontaneously develop PAH. These findings support the rationale for exploring dysregulated polyamine homeostasis in PAH and suggest a potential for therapeutic targeting of polyamine pathways in PAH.

## 1. INTRODUCTION

Pulmonary arterial hypertension (PAH) is a progressive vascular disorder characterised by the narrowing and obliteration of small precapillary lung arterioles ^1^. Endothelial cell (EC) dysfunction, proliferation of mesenchymal cells in the vascular wall and aberrant inflammation ^1, 2^ contribute to this pathological process. Despite the availability of licensed therapies, PAH patient survival remains poor, necessitating new targeted treatments.

The identification of heterozygous germline mutations in the bone morphogenetic protein type II receptor (*BMPR2*) gene ^3, 4^ and more recent identification of loss-of-function mutations in other BMP pathway components ^5^ has underpinned potential PAH therapies to enhance BMP signalling ^6^. However, some rare PAH-related genes appear distinct from the BMP pathway ^5, 7^, suggesting additional mechanisms underlying the pathobiology of PAH that may be informative for alternative therapies.

In a European-wide PAH cohort study, we identified 11 rare heterozygous *ATP13A3* variants with protein-truncating variants overrepresented (6 of 11), suggesting a loss-of-function in PAH^7^. Since then, more *ATP13A3* variants have been reported in other PAH patient cohorts ^8-10^. Though ATP13A3 is expressed in various cell types, including pulmonary vascular cells, pulmonary macrophages and dendritic cells ^7, 11, 12^, its function remains unclear. *ATP13A3* is a member of the P5B-type ATPase family (ATP13A2-5) and ATP13A2 has recently been identified as a polyamine transporter^13^. As ATP13A3 has close homology to ATP13A2 and *ATP13A3* mutations account for the polyamine uptake deficiency in CHO-MG cells ^14^, ATP is thought to be a polyamine transporter.

Cellular polyamine levels are tightly regulated through the integration of their biosynthesis/catabolism and their transport, and disruption of these pathways can lead to diseases ^15, 16^. Here we established that ATP13A3 mediates cellular polyamine uptake in human vascular ECs, whereas PAH-associated ATP13A3 mutations reduce polyamine transport. Loss of ATP13A3 leads to PAH-associated phenotypes in pulmonary arterial ECs and in mice harbouring a PAH-associated *Atp13a3* frameshift variant (P452Lfs). Collectively, our data explain the impact of ATP13A3 mutations in PAH and suggest that dysregulated polyamine homeostasis may contribute to its pathobiology.

## 2. Methods

Key protocols are described here and additional detailed protocols are described in the online data Supplement.

### 2.1. Animals

All animal procedures were performed in accordance with the Home Office Animals (Scientific Procedures) Act (1986) and were approved under Home Office Project Licence 70/8850. The *Atp13a3* genetically modified mouse, designated *Atp13a3*^P452Lfs^, was generated at MRC Harwell using CRISPR-Cas9 editing to introduce a 1nt deletion resulting in a frameshift and termination codon after a further 7 amino acids (P452LfsTer7), equivalent to the human PAH variant P456Lfs ^7^. The mice were viable, fertile, do not exhibit any behavioural abnormalities and do not differ in appearance or weight from their wild-type littermates.

### 2.2. Haemodynamic assessments of mice

Cardiac catheterization was performed via the closed-chest technique, as described previously ^17^. Measurements of RV systolic pressure (RVSP) were performed under isoflurane anaesthesia (2.0-2.5% isoflurane, 100% oxygen 2 L/min) in spontaneously breathing animals. In the same animals, systolic (SBP) in the aorta was measured.

Hearts were excised and the right ventricle (RV) dissected from the left ventricle plus septum, weighed and then fixed in 10% neutral-buffered formalin. The lungs were inflated with 10% neutral-buffered formalin and harvested for histological analyses as described in the online supplement.

Echocardiographic ultrasound measurements of heart rate, right ventricular dimensions, and PA pressure surrogates were conducted in spontaneously breathing animals under isoflurane anesthesia using an ultrasound machine (Vevo 3100 System, VisualSonics Inc.) equipped with a 40MHz linear array transducer.

### 2.3. Cell culture

Human pulmonary artery endothelial cells (hPAECs) were purchased from Promocell (Heidelberg, Germany) and maintained in EGM2 media (Promocell) with 2% FBS and antibiotic/antimycotic, as per the supplier’s instructions.

Human blood outgrowth ECs (BOECs) were isolated from 40-80ml of blood as previously described ^18^. BOEC lines were grown in EGM-2 with the addition of 10% FBS, antibiotic/antimycotic and omission of heparin. For experiments involving BOEC generation, all donors provided informed written consent under human study 07/H0306/134 (Cambridgeshire 3 Research Ethics Committee) or REC - 17/LO/0563 (ATP13A3-LK726X variant carrier). Demographic and mutation information for the BOECs used in this study is specified in supplement table 1. Both hPAECs and BOECs were used for experiments between passage 4 to 7.

The immortalised (SV40-transformed) human microvascular endothelial cell-1 (HMEC-1) line was purchased from ATCC. HMEC-1 were grown in MCDB131 medium (without glutamine) (Thermo Fisher Scientific) supplemented with 1 µg/ml Hydrocortisone (Sigma), 10 mM Glutamine (Sigma), 10 ng/ml Epidermal Growth Factor (EGF) (R&D systems), 10%(v/v) FBS and antibiotic/antimycotic. All cells were routinely tested for mycoplasma and were only used if negative.

### 2.4. Cellular transfection and transduction

Cells were transfected with *ATP13A3* siRNA, *ATP13A3* expression plasmids or transduced with lentiviral expression particles ^19^ as described in the online data Supplement.

### 2.5. Measurement of cellular polyamines

Aqueous metabolites were extracted from cell lysates and analysed for polyamine content by LC-MS as described in the online supplement.

### 2.6. BODIPY-Labelled polyamine uptake assay

Spermine-BODIPY, Spermidine-BODIPY, and Putrescine-BODIPY were synthesized as previously described ^20^ and dissolved in 0.1 M MOPS, pH = 7.0 (AppliChem, A1076). Uptake of Polyamine-BODIPY in HMEC-1 was determined by flow cytometry. HMEC-1 cells were seeded in 12-well plates at 300,000 cells/well and left to attach overnight. After determining the kinetics of uptake to ensure being in the linear phase, the cells were then incubated with the respective polyamine-BODIPY concentration (5 µM if single concentration) for 30 min after which they were trypsinised, centrifuged at 300 x g and the pellet washed with cold Dulbecco’s Phosphate Buffered Saline (PBS) solution without calcium or magnesium (Sigma, D8537). The pellets were then re-suspended in 1% BSA/PBS. Polyamine-BODIPY uptake was determined by flow cytometry on a BD FACSCanto™ II instrument, with 10,000 events recorded per treatment.

### 2.7. Statistical analysis

Data are presented as mean ± S.E.M and are analysed using GraphPad Prism 7. All data shown is n=3 (unless mentioned otherwise) where n represents the number of independent repeats. Kolmogorov-Smirnov test was performed for assessing normality and data were analysed by one-way analysis of variance with post hoc Tukey’s HSD analysis, One-/Two-way ANOVA followed by multiple comparisons using Dunnett’s/Tukey’s post hoc tests, or unpaired two-tailed student’s t-test as indicated. P<0.05 is considered statistical significance.

## 3. RESULTS

### 3.1. ATP13A3 is a polyamine transporter localised to recycling endosomes

P5B-ATPases are multi-span transmembrane proteins that may localise to the plasma membrane ^21^ or endosomal system ^22^. Endogenous ATP13A3 in human pulmonary artery ECs (hPAECs) localised primarily to a perinuclear region with lower nuclear staining (figure 1A). Both GFP-tagged (figure 1B-C) and endogenous (supplement figure 1) ATP13A3 in HMEC-1 cells co-localised mainly with Rab11, suggesting ATP13A3 resides primarily in recycling endosomes in ECs.

**Figure 1.**
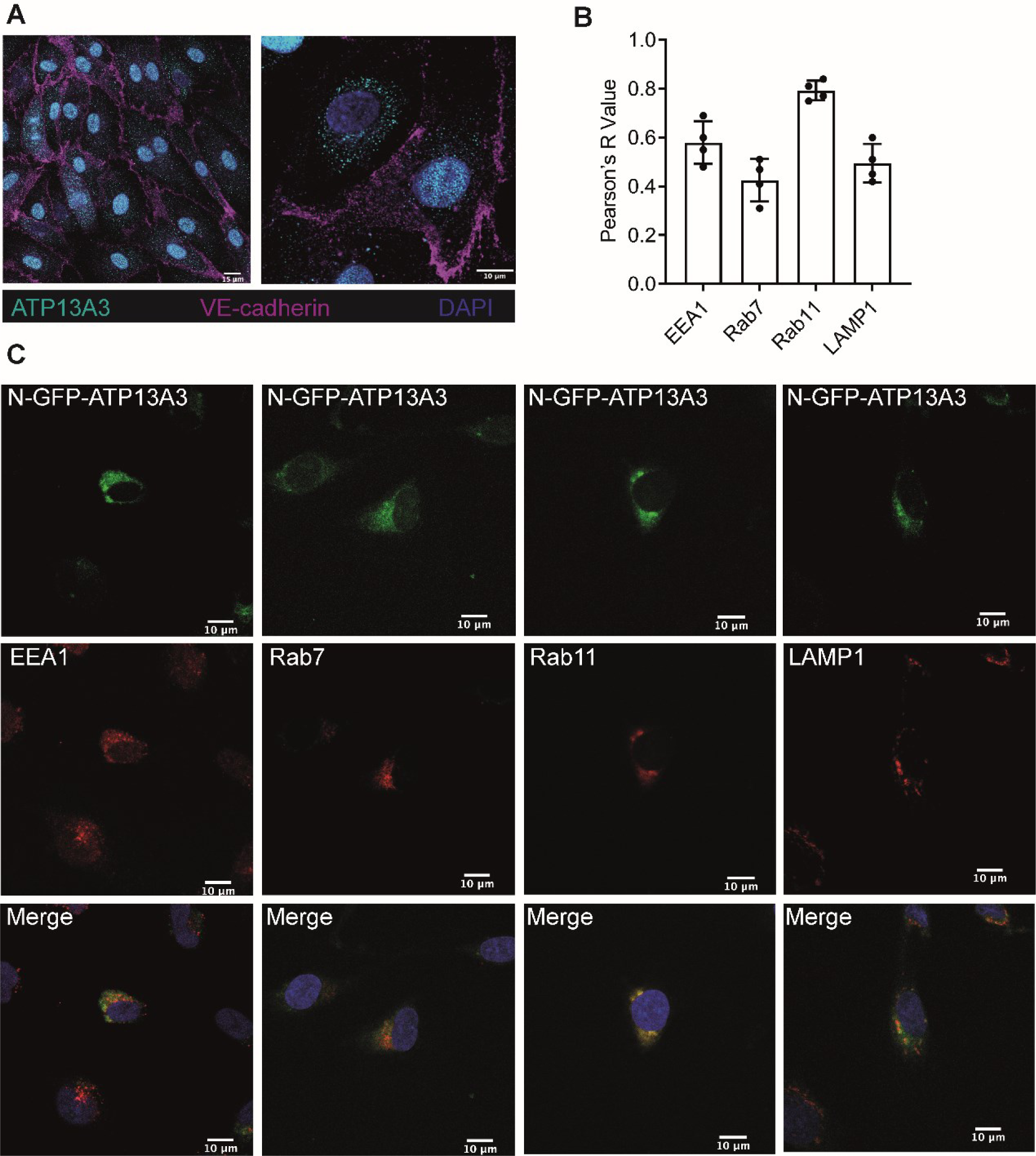
ATP13A3 is a polyamine transporter residing in the recycling endosome of endothelial cells. (A) Confocal images at 40X (left panel, scale bar = 15µm) and 63X (right panel, scale bar = 10 µm) of hPAECs co-stained with anti-ATP13A3 and anti-VE-Cadherin. (B) Pearson’s coefficients of the correlation of GFP-tagged ATP13A3 to different endosomal markers in HMEC-1 cells. (C) Confocal images (63X, scale bar = 10 µm) of HMEC-1 cells transiently overexpressing hATP13A3-N-GFP-pcDNA6.2 co-stained with antibodies against either EEA1, Rab7, Rab11 or LAMP1. Data are representative of n=4 experiments.

We hypothesised that ATP13A3 mediates polyamine transport in primary human ECs. *ATP13A3* siRNA (si*ATP13A3*) in hPAECs reduced *ATP13A3* expression without affecting the expression of *ATP13A1-2* (supplement figure 2A-C). We could not detect *ATP13A4-5* expression in these cells. si*ATP13A3* significantly reduced the basal cellular putrescine, spermine and spermidine contents in hPAECs, while only the increase in putrescine content was significantly attenuated upon exogenous polyamine supplementation (figure 2A). For validation, we stably silenced *ATP13A3* in HMEC-1 cells (HMEC-1^mi*ATP13A3*^) using three lentiviral micro-RNAs (miRNA) targeting different regions of the *ATP13A3* mRNA (figure 2B and supplement figure 2D-E). Like in hPAECs, basal putrescine, spermidine and spermine contents were reduced in HMEC-1^mi*ATP13A3*^ cells (supplement figure 2F) and uptake of BODIPY-tagged putrescine (PUT-BDP) and Spermidine (SPD-BDP), but not Spermine (SPM-BDP) was impaired (figure 2C). Furthermore, confocal imaging confirmed lower PUT-BDP uptake in HMEC-1^mi*ATP13A3*^, with the internalized polyamine predominantly confined to punctae (figure 2D). In conclusion, ATP13A3 determines polyamine uptake, redistribution and homeostasis in ECs.

**Figure 2.**
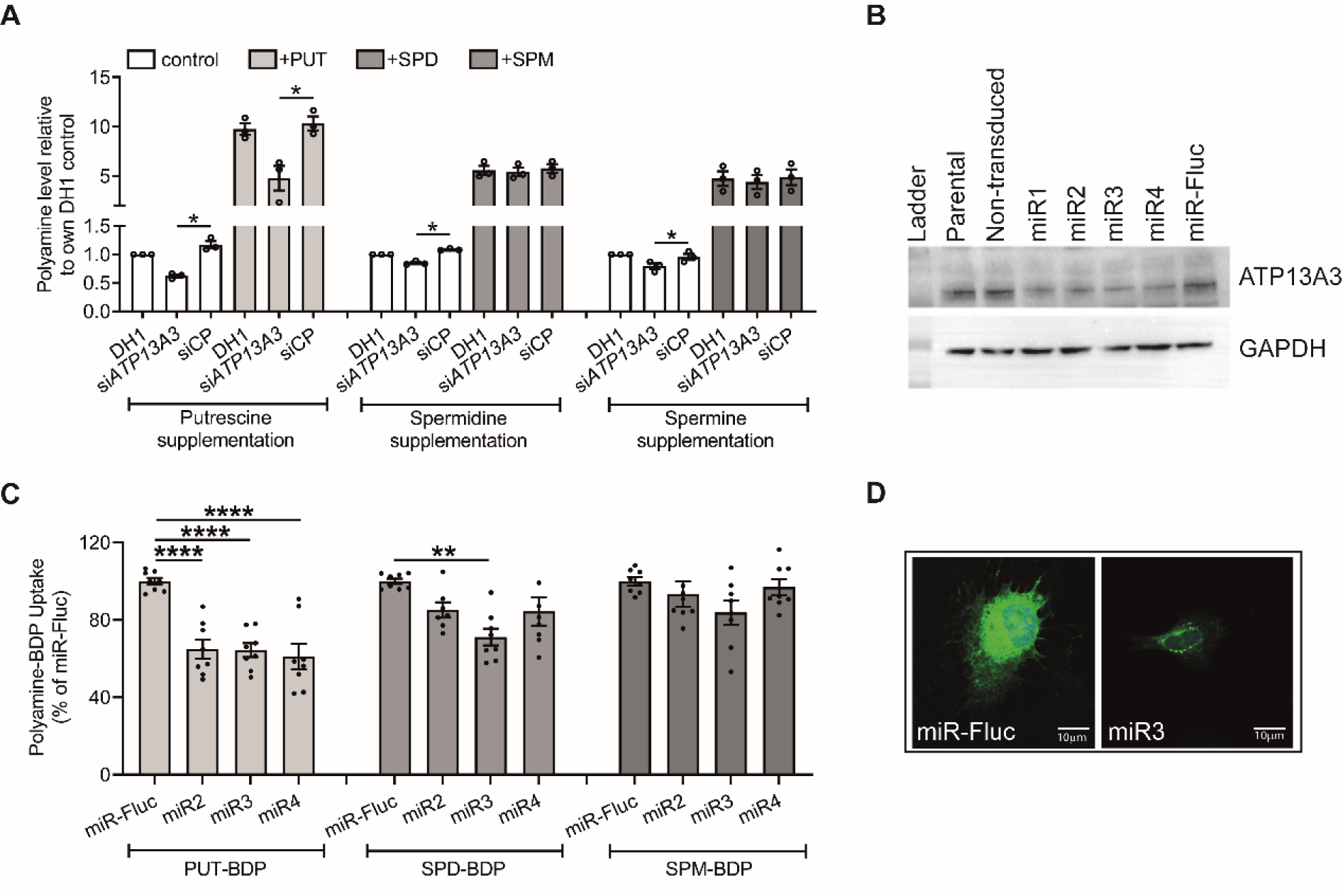
ATP13A3 deficiency impairs polyamine transport in endothelial cells. (A) Cellular putrescine (PUT), spermidine (SPD) and spermine (SPM) levels in hPAECs measured by LC-MS. Cells were transfected with DharmaFECT1™ (DH1) alone, si*ATP13A3* or non-targeting siRNA control (siCP) and cultured overnight in EBM2 containing 2% FBS supplemented with or without 1mM putrescine, 10µM spermidine, or 10µM spermine. Data (n=3 experiments) are presented as polyamine peak area ratio relative to 2% FBS DH1. (B) Western Blot showing ATP13A3 protein expression in Parental, Non-transduced, and HMEC-1 cells stably expressing miRNAs targeting ATP13A3 (miR1–miR4), with miRFLUC (Firefly Luciferase) as a control. (C) BODIPY (BDP)-labelled polyamine uptake in HMEC-1 stable knockdown lines (n=4 experiments, two technical replicates per experiment). Data are normalised to the mean fluorescent intensities of miR-Fluc. (D) Confocal microscopy depicting the uptake and distribution of PUT-BDP in HMEC-1 cells expressing miRFLUC and ATP13A3 miR3 following 2 h treatment with PUT-BDP (scale bar = 10 µm). (A,C) Data (mean ± SEM) were analysed using a one-way ANOVA with Tukey’s post hoc test for multiple comparisons *P<0.05 **P<0.01, ****P<0.0001.

### 3.2. ATP13A3 levels alter the expression of polyamine biosynthesis pathways

Polyamine homeostasis is maintained through the integrated functions of polyamine transporters and polyamine metabolism enzymes (supplement figure 3). Ornithine decarboxylase (ODC) converts ornithine into putrescine to initiate polyamine biosynthesis and is tightly regulated by cellular polyamine levels ^16^. ODC protein levels increased without altering mRNA expression in si*ATP13A3* transfected hPAECs or BOECs (figure 3A-B and supplement figures 4 and 5A-C). Moreover, expression of antizyme (OAZ), which mediates the polyamine-dependent proteasomal degradation of ODC ^23^ was reduced, whereas the antizyme inhibitor (AZIN) was unchanged (figure 3B and supplement figure 5C). Hence, the increased ODC protein in *ATP13A3* deficiency may occur via *OAZ* downregulation.

**Figure 3.**
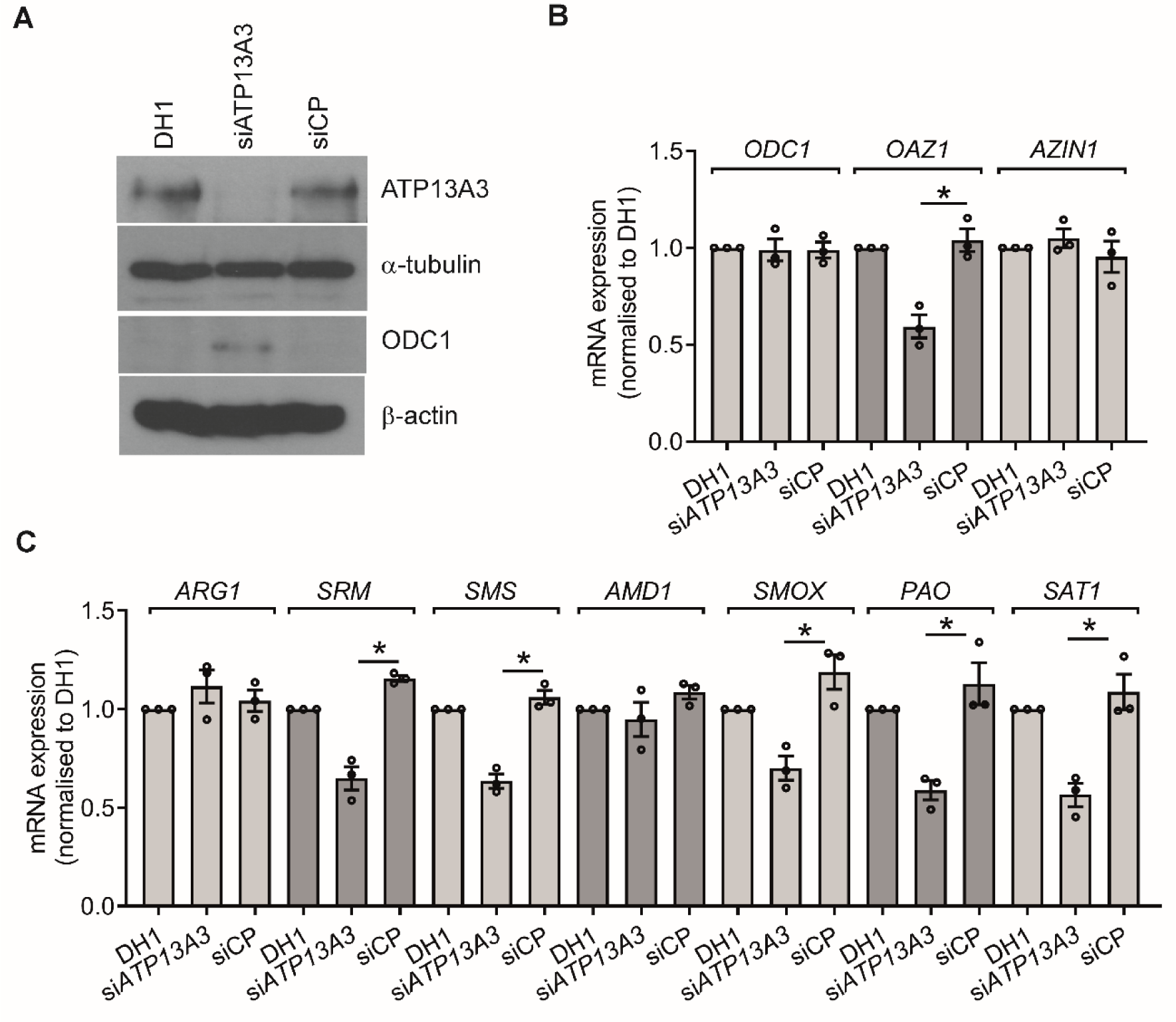
ATP13A3 deficiency affects polyamine metabolism in hPAECs. (A) Immunoblotting for ATP13A3 and ODC1 in hPAECs after transfection with DharmaFECT1™ (DH1) alone, si*ATP13A3* or non-targeting siRNA control (siCP). (B,C) Transcription of (B) *ODC1, OAZ1, AZIN1*, polyamine biosynthesis enzymes (*ARG1, SRM, SMS, AMD1*) and (C) catabolic enzymes (*SMOX, PAO, SAT1*) in hPAECs transfected with DH1, si*ATP13A3* or siCP. Data (n=4 experiments) are fold-change relative the DH1 control for each transcript. (B,C) Data (mean ± SEM) were analysed using a One-way ANOVA with Tukey’s post hoc test for multiple comparisons. *P<0.05, **P<0.01 compared to siCP.

In si*ATP13A3*-transfected hPAECs and BOECs, arginase 1 (*ARG1*) and adenosylmethionine decarboxylase (*AMD1)* expression were unchanged, whereas spermidine synthase (*SRM*) and spermine synthase (*SMS*) were reduced (figure 3C and supplement figure 5D). Also, the expression of the polyamine catabolic enzymes, spermidine/spermine N1-acetyltransferase 1 (*SAT1*) and polyamine oxidase (*PAO*) were lower in si*ATP13A3*-transfected cells, possibly as an attempt by the cells to rebalance putrescine levels (figure 3C and supplement figure 5D).

### 3.3. ATP13A3 deficiency leads to pulmonary artery endothelial dysfunction

Dysregulated proliferation and increased apoptosis and permeability of ECs contribute to the pathobiology of PAH ^1, 2^. We previously reported that *ATP13A3* knockdown in BOECs impaired their proliferation and increased apoptosis in reduced serum ^7^. Here, we established that si*ATP13A3* reduced hPAEC proliferation (figure 4A), which was associated with reduced mRNA expression of cyclins E (*CCNE1*), A (*CCNA1*), and B (*CCNB1*), suggesting impaired cell cycle progression (figure 4B). Supplementation with 10 µM putrescine, spermidine or spermine promoted hPAEC proliferation (supplement figure 6A-C), which was attenuated in si*ATP13A3*-transfected hPAECs (supplement figure 6D). The reduced proliferation in EBM2/2% FBS was not due to increased apoptosis as caspase 3/7 activity was not altered by ATP13A3 knockdown (figure 4C). However, caspase 3/7 activity was increased by si*ATP13A3* in hPAECs incubated in low serum conditions, suggesting a greater susceptibility to intrinsic stress (figure 4C). Moreover, si*ATP13A3* did not alter basal hPAEC monolayer permeability, but we observed 40% higher permeability when monolayers were exposed to 1 U/ml thrombin (figure 4D).

**Figure 4.**
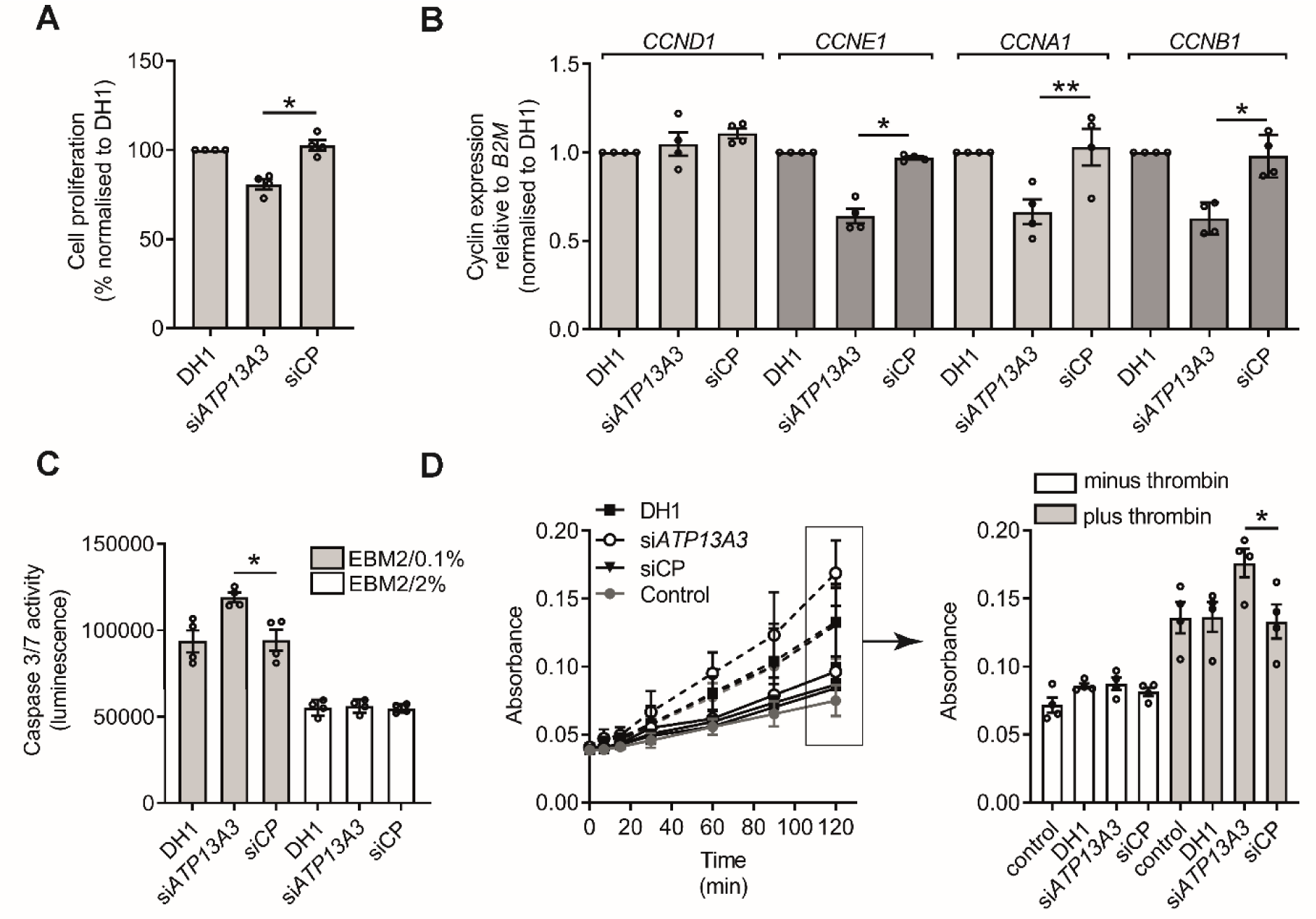
*ATP13A3* deficiency leads to endothelial dysfunction. (A) Proliferation of transfected hPAECs over 6 days in EBM2 with 2% FBS, with media replenished every 2 days. (B) Transcription of *CCND, CCNE, CCNA, CCNB* mRNAs with *ATP13A3* deficiency assessed by qPCR. Data are fold-change relative to the DhamaFECT1™ (DH1) control for each transcript. (C) Apoptosis assessed by Caspase-Glo®3/7 assay of transfected hPAECs cultured in EBM2 with 0.1% FBS or 2% FBS. (D) Permeability of transfected hPAEC monolayers to horseradish peroxidase in the absence or presence of 1U/ml Thrombin assessed by colorimetric assay. The left panel shows the time-course and the right panel shows the raw absorbance values for the different groups at the 2 h time point. Data (n=4 experiments) in panels A-D are mean ± SEM and were analysed using a One-way ANOVA with Tukey’s *post hoc* test for multiple comparisons. *P<0.05 **P<0.01 compared to siCP.

### 3.4. PAH associated mutations impair ATP13A3-mediated putrescine uptake in ECs

To establish if PAH-associated *ATP13A3* variants are pathogenic, we assessed the functional impact of five PAH-associated missense variants (L675V, M850I, V855M, R858H, L956P; supplement figure 7A) in different EC models, comparing these to ATP13A3-wild-type (ATP13A3-WT) protein and a transport dead mutant with a D498N substitution in the catalytic autophosphorylation domain.

Stable lentiviral overexpression of the untagged WT, but not the D498N mutant, increased the uptake of PUT-BDP and SPD-BDP, but not SPM-BDP (figure. 5A-B and supplement figure 7B), consistent with our knockdown results (figure 2C). Benzyl Viologen (BV), a polyamine uptake inhibitor ^14^, blocked PUT-BDP uptake (supplementary figure 7C).

Interestingly, despite their mRNA expression levels being comparable to the WT (supplement figure 7E), two disease variants (V855M and R858H) showed reduced protein expression (figure 5C and supplement figure 7F). Since the antibody epitope (488-631) lies outside the mutated region, this implies reduced protein stability. Consequently, these variants fail to increase PUT-BDP and SPD-BDP uptake (figure 5A-B). Though the other variants expressed well, the uptake of both PUT-BDP and SPD-BDP was impaired only for the L956P variant. Intriguingly, the L675V variant only demonstrated reduced SPD-BDP uptake, suggesting altered substrate specificity.

**Figure 5.**
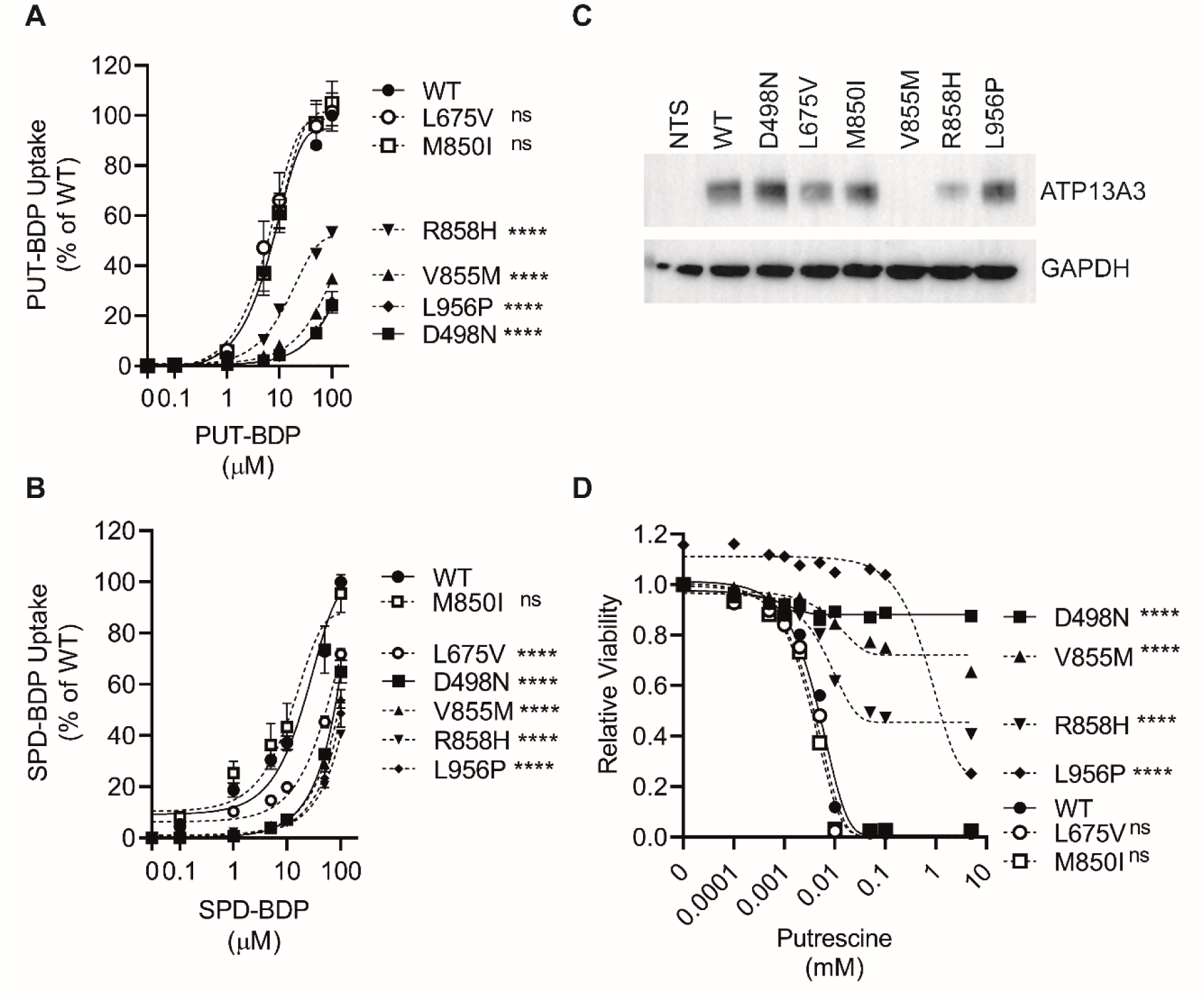
PAH-associated *ATP13A3* variants exhibit deficient polyamine uptake. (A,B) Flow-cytometry analysis for assessment of cellular uptake of increasing concentrations of (A) PUT-BDP (n=5 experiments) and (B) SPD-BDP (n=6 experiments) after 30 minutes exposure. Data are normalised to WT. (C) Western blot showing ATP13A3 protein expression in non-transduced (NTS) HMEC-1 cells compared to those stably expressing untagged ATP13A3 WT (WT), a transport dead mutant (D498N) or five PAH-associated variants (L675V, M850I, V855M, R858H and L956P). (D) Cytotoxicity (MUH reagent) assay with a concentration-response analysis to putrescine PUT (N=6 experiments). (A, B, D) Data were analysed by two-way ANOVA followed by multiple comparisons using Tukey’s post hoc tests. **** p<0.00005 and ns = non-significant.

Polyamines are essential for cell survival but at high concentrations, can be cytotoxic ^13^. Stable ATP13A3-WT overexpression sensitised HMEC-1 cells to escalating putrescine concentrations whereas the D498N mutant did not (figure 5D). Mirroring their impact on PUT-BDP uptake, the three untagged PAH-associated variants (V855M, R858H and L956P) also exhibited reduced sensitivity to putrescine toxicity.

So far, the M850I variant did not differ from WT, although the endogenous putrescine, spermidine and spermine levels were reduced in HMEC-1 cells stably over-expressing the M850I, L956P variants and D498N mutant, albeit non-significantly (supplement figure 8A-C). However, stable overexpression of the L675V and M850I mutations did significantly decrease basal spermidine and spermine levels in neuroblastoma SH-SY5Y cells (supplement figure 8D-F), suggesting a cell-type specific phenotype for this mutation.

To analyse the intracellular localization of the variants, we transiently overexpressed GFP-tagged WT, D498N and the fiver PAH-associated ATP13A3 variants in HMEC-1 cells. Surprisingly, all GFP-tagged variants and WT-GFP consistently colocalised with Rab11, suggesting that when transiently expressed, V855M-GFP and R858H-GFP express well (supplement figure 9A). In hPAECs transiently overexpressing these constructs, basal putrescine levels were similar with all variants (supplement figure 9B). Supplementation with 1 mM putrescine increased endogenous putrescine levels mainly in WT-GFP expressing cells, since the ATP13A3 variants attenuated (R858H-GFP and L956P-GFP) or abolished (D498N-GFP, L675V-GFP, M850I-GFP and V855M-GFP) this response (supplement figure 10C). We confirmed similar *ATP13A1-3* expression of all the constructs (supplement figure 10A-C).

We further assessed the impact of high polyamine concentrations on apoptosis (caspase-3/7 activity). Although ATP13A3-WT-GFP overexpression sensitised hPAECs to 10 mM putrescine (supplement figure 11A), this was reduced for the D498N-GFP mutant and disease variants L675V-GFP, M850I-GFP and V855M-GFP, consistent with the attenuated response to putrescine supplementation in these cells. In contrast, the R858H-GFP and L956P-GFP variants behaved more akin to the ATP13A3-WT (supplement figure 11A). No differences were seen between ATP13A3-WT, the D498N and PAH related variants to spermidine and spermine toxicity (supplement figure 11B-C).

Together, our analysis in complementary expression systems reveals that the ATP13A3 missense variants present different forms of loss-of-function phenotypes affecting polyamine uptake and/or homeostasis.

### 3.5. The ATP13A3^LK726^ frameshift mutation predisposes BOECs to apoptosis by affecting ATP13A3-mediated polyamine transport

To cross-validate our findings in a disease-relevant endogenous system, we derived BOECs from a PAH patient bearing a heterozygous *ATP13A3* frameshift variant (*ATP13A3*^LK726X^, c.2176_2180delTTAAA), confirmed by Sanger sequencing (supplement figure 12). This mutation creates a premature stop codon (TGA 733X), predicted to reduce *ATP13A3* mRNA expression through nonsense-mediated decay (supplement figure 12B). Indeed, both *ATP13A3* mRNA and protein levels (figure 6A and supplement figure 13A) were reduced in *ATP13A3*^LK726X^ BOECs compared to control BOECs without changes in *ATP13A1* and *ATP13A2* (supplement figure 13B-C). Compared to control cells, *ATP13A3*^LK726X^ BOECs exhibited lower basal putrescine content, which failed to increase upon exogenous putrescine supplementation (figure 6B). Although basal and supplemented spermine and spermine contents were unchanged (figure 6B), the uptake of PUT-BDP and SPM-BDP were significantly lower than in control BOECs (figure 6C). Functionally, caspase-3/7 activity in low serum was significantly elevated in *ATP13A3*^LK726X^ BOECs (figure 6D) and this was partially rescued by overexpression of the ATP13A3-WT, but not the D498N mutant (supplement figure 14).

**Figure 6.**
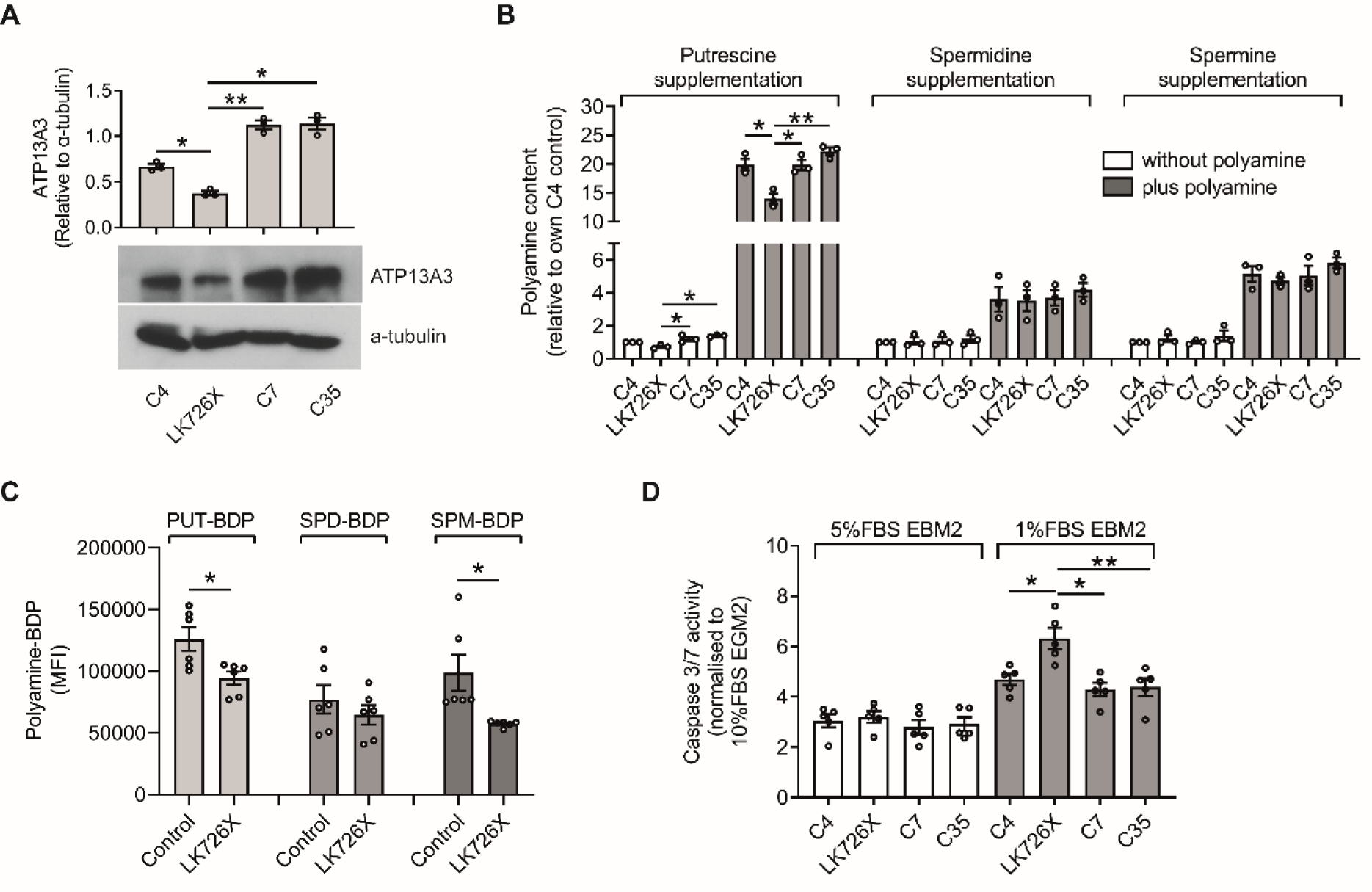
The *ATP13A3* LK726X frameshift mutation predisposes BOECs to apoptosis by affecting ATP13A3-mediated polyamine transport. (A) Immunoblotting of ATP13A3 in control BOECs (C4, C7, C35) and *ATP13A3*^LK726X^ BOECs. Densitometric analysis of ATP13A3 and α-tubulin was performed (Graph). (B) Cellular polyamine contents, measured by LC-MS, of BOECs in media or supplemented with 1mM putrescine, 10µM spermidine or 10µM spermine. Data (n=3 experiments) are presented as polyamine peak area ratio relative to the C4 control BOEC line. (C) BOEC uptake of PUT-BDP, SPD-BDP and SPM-BDP measured by flow cytometry (n=6 experiments). (D) Cell apoptosis of BOECs cultured in EBM2 supplemented with 1% FBS or 5% FBS was assessed by Caspase-Glo®3/7 assay (n=5 experiments). Data are normalised to cells cultured in EGM2 containing 10% FBS. (A-D) Data are mean ± SEM analysed using (A,B,D) one-way ANOVA with Tukey’s post hoc test for multiple comparisons or (C) unpaired t-test. *P<0.05 **P<0.01 compared with *ATP13A3*^LK726X^.

Interestingly, *ODC* mRNA and protein levels were increased in *ATP13A3*^LK726X^ BOECs (supplement figure S15A-B), which was not explained by *OAZ1* changes, but most likely by higher *AZIN1* expression (supplement figure S15C-D). Intriguingly, expression of the synthetic enzymes, *ARG1*, *AMD1*, *SRM* and *SMS* were also elevated (supplement figure S15E), while the catabolic enzymes remained comparable between *ATP13A3*^LK726X^ and control BOECs (supplement figure S16). Collectively, our data suggest the *ATP13A3*^LK726X^ mutation disrupts polyamine homeostasis in BOECs, predisposing cells to apoptosis.

### 3.6. Mice harboring an Atp13a3^P452Lfs^ mutation spontaneously develop PAH

We observed that animals heterozygous for an *Atp13a3* frameshift variant (P452Lfs), previously identified in patients with PAH [7], exhibit reduced lung *Atp13a3* expression (figure 7A). Heterozygous *Atp13a3*^P452Lfs^ mice spontaneously and consistently developed pulmonary hypertension compared to their littermate controls, with a significant elevation of right ventricular systolic pressure (figure 7B) while heart rate and systemic blood pressure did not differ (supplement figure 17A-B). Transthoracic echocardiography demonstrated a significant shortening of pulmonary artery acceleration time (PAAT, a surrogate of pulmonary artery pressure and pulmonary vascular resistance) in heterozygous *Atp13a3*^P452Lfs^ mice (supplement figure 17C). Heterozygous *Atp13a3*^P452Lfs^ mice also exhibited increased right heart dimensions, namely right ventricular inner diameter (supplement figure 17D) and right ventricular end diastolic anterior wall thickness (supplement figure 17E) compared to the wild-type littermates. Again, heart rate did not differ between the two groups (supplement Figure 17F).

**Figure 7.**
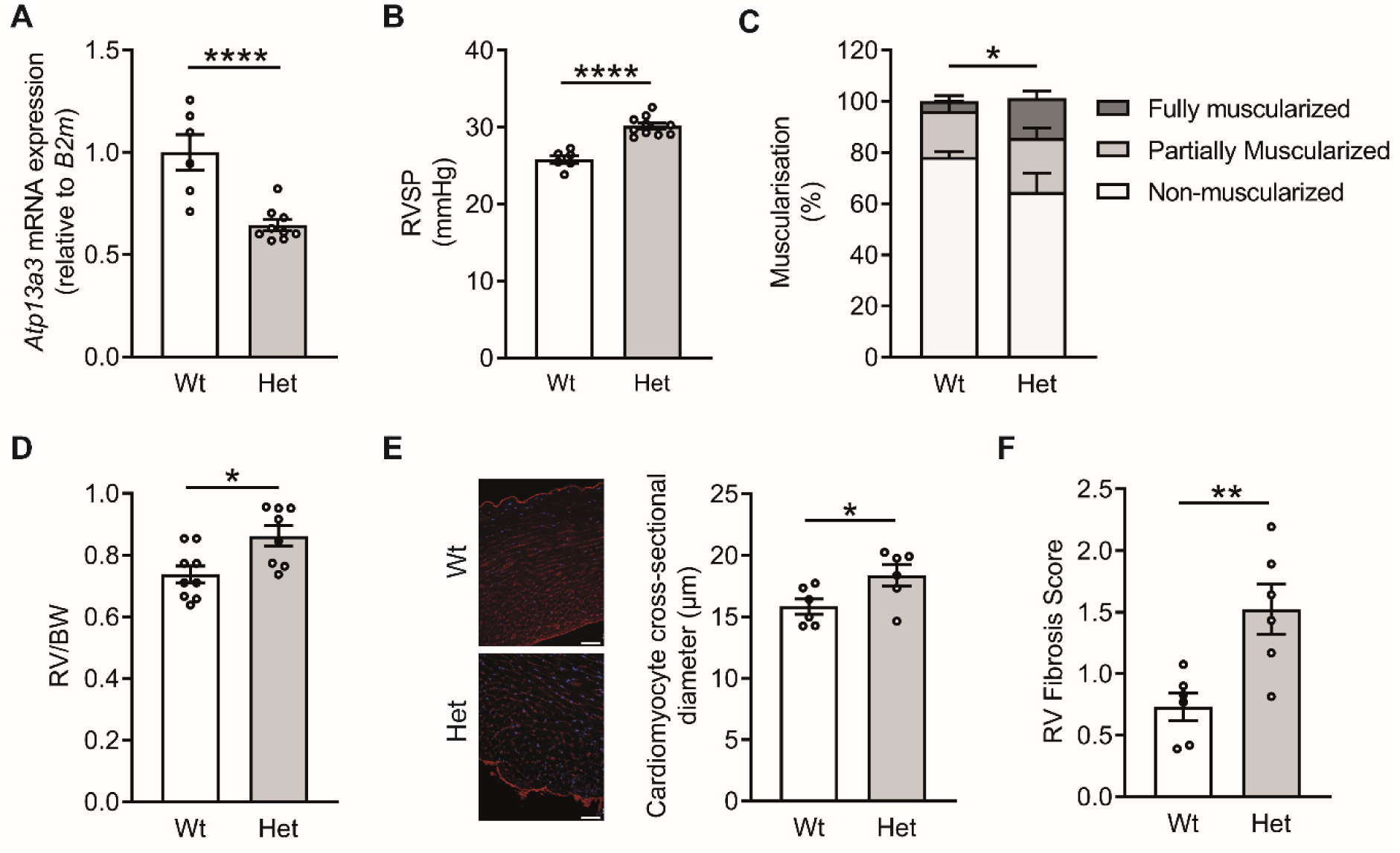
Mice harbouring an *Atp13a3*^P452Lfs^ variant spontaneously develop PAH. (A) Expression of *Atp13a3* mRNA in whole lungs of *Atp13a3*^P452Lfs^ heterozygous mice and controls (n = 6, 9) (B) Invasive hemodynamic measurement of right ventricular systolic pressure (RVSP) in *Atp13a3*^P452Lfs^ heterozygous mice and controls (n=6, 10). (C) Quantification of vessel muscularisation in arterioles <50 μm in diameter (n=5 mice per group, n = 4-8 to 10 vessels/lung). (D) Ratio of right ventricular (RV) weight to bodyweight (BW) (n=9, 8). (E) Wheat-germ agglutinin staining of cardiomyocytes in heart sections (scale bar = 50µm) from which the cardiomyocyte cross-sectional areas were measured (*graph*, n=6 per group). (F) Scoring for right ventricular fibrosis in picrosirius red-stained sections (n=6 per group). Data are mean ± SEM analysed by a two-tailed t-test with Welch’s correction, *P<0.05, ****P<0.0001.

Consistent with the haemodynamic data suggesting a baseline PAH phenotype, we also observed histological changes in the pulmonary circulation and hearts of *Atp13a3*^P452Lfs^ mice. Lung vascular morphometric analysis revealed that heterozygous *Atp13a3*^P452Lfs^ mice had a significantly higher percentage of fully-muscularized small pulmonary arteries (figure 7C). Consistent with this phenotype, the right ventricles were also hypertrophic (figure 7D). This was associated with a significant increase in cardiomyocyte cross-sectional diameter (figure 7E) and in interstitial fibrosis (figure 7F). Although milder than the PAH observed in induced models of disease, these data confirmed that *Atp13a3*^P452Lfs^ heterozygous mice spontaneously and consistently demonstrated a PAH phenotype without the need for an applied disease-promoting stimulus.

## 4. DISCUSSION

In this study we establish ATP13A3 in ECs as a recycling endosome polyamine transporter enabling cellular polyamine uptake, the deficiency of which leads to EC dysfunction. Human disease-associated ATP13A3 variants present a loss of polyamine uptake phenotype, whereas mice harbouring a human disease relevant *Atp13a3* mutation develop PAH. This suggests that disrupted polyamine homeostasis represents a new genetic mechanism promoting PAH pathobiology.

ATP13A3 most likely functions as a regulator of cellular polyamine content from recycling endosomes ^14, 21, 24^. This function mirrors the closely-related ATP13A2, a late endo/lysosomal polyamine exporter that delivers polyamines into the cytosol, preventing lysosomal accumulation while regulating cellular content ^13^. We studied several EC models of ATP13A3 deficiency, namely transient siRNA, stable miRNA and patient-derived *ATP13A3*^LK726X^ BOECs in parallel to overexpression experiments. We found that *ATP13A3* deficiency in these EC models suppressed the uptake of BDP-labelled polyamines. The reduction in uptake of each polyamine differed between the cell models, possibly reflecting model-specific variations in the receptor-mediated endocytosis pathway proximal to ATP13A3-mediated transport to the cytosol ^16^. We also observed reductions in endogenous polyamine levels in ATP13A3 deficient EC cell models, and an attenuated response to putrescine supplementation. Conversely, ATP13A3-WT over-expression led to increased uptake of mainly PUT-BDP and SPD-BDP, and a higher intracellular putrescine content. The differences observed between labelled polyamine uptake versus endogenous levels are not unusual. Fluorescently labelled polyamines behave similarly to radiolabelled polyamines, thus directly representing uptake ^24^. In contrast, quantifying endogenous polyamine levels by mass-spectrometry reflects an integration of the polyamine metabolic pathways, so the uptake of one polyamine species may impact all the polyamine species.

In this study, we have also identified that the loss of ATP13A3 not only reduces intracellular polyamine levels but also leads to a reprogramming of the polyamine metabolism pathway. Despite subtle differences between our EC models, we generally observed an increased polyamine biosynthesis (*e.g.* upregulated *ODC1/AZIN1* or reduced *OAZ1*) as well as a reduced polyamine catabolism (*e.g.* lower SAT1 and/or PAO), both of which may compensate for polyamine loss. This probably explains why multiple endogenous polyamine species are simultaneously affected. One limitation of cells isolated from PAH patients is the potential impact of changes acquired due to the disease state ^25^ and the heterozygous expression of the variant. These may explain why transcript alterations of polyamine biosynthesis enzymes with short term *ATP13A3* loss (siRNA transfection) differed from chronic deficiency (genetic defects).

In this context of altered polyamine metabolism, it remains difficult to deduce the precise polyamine transport specificity of ATP13A3 from cellular data. Our observations suggest that ATP13A3 displays a broad polyamine selectivity that exerts the strongest impact on putrescine uptake and content. Similar findings were also reported for Atp13a3 in CHO-MG cells, where unlabelled spermidine and spermine competed with ATP13A3-dependent PUT-BDP uptake ^14^.

The dual effect of *ATP13A3* deficiency on polyamine uptake and polyamine homeostasis in ECs may explain the strong impact of ATP13A3 on endothelial health and functionality. Polyamines are essential for cell growth ^15, 16^, with polyamine depletion leading to cell cycle arrest ^16^. *ATP13A3* deficiency suppressed serum-dependent proliferation and *cyclin A, E and B* mRNA expression in BOECs implying repression of G1-S transition and DNA synthesis in the cell cycle. Polyamine depletion may promote cell apoptosis in response to pathogenic insults ^26, 27^. We show that *ATP13A3* deficiency increases apoptosis in hPAECs, BOECs and *ATP13A3*^LK726X^ BOECs when they are stressed by serum starvation, an effect that is rescued with ATP13A3-WT overexpression. Moreover, polyamines are essential for epithelial cell-cell junctions ^28^ and we observed that *ATP13A3* deficiency exacerbated thrombin-dependent hPAEC monolayer permeability, indicating an important role of *ATP13A3* in maintaining endothelial integrity.

Using our EC models, we examined the functional impact of PAH-associated missense variants using lentiviral overexpression. A homozygous V855M variant was identified in a child with early-onset of PAH leading to early death ^29^. The other PAH missense variant (L675V, M850I, R858H and L956P) were heterozygous and found in older PAH patients ^7^. Collectively, our data confirm pathogenicity due to loss-of-function, though possibly via differing impacts on ATP13A3 function, negatively impacting on (i) protein expression, (ii) transport activity, (iii) polyamine homeostasis and/or (iv) substrate specificity. These differences may relate to cell type (*e.g.* for M850I), transient *versus* the lentiviral stable mutant expression (the latter being more potent potentially increasing the window), and/or the possible stabilizing effect of the GFP tag (for V855M and R858H). The latter may explain the low protein stability of untagged V855M and R858H variants compared to GFP fusion products. Furthermore, the variable impacts of these ATP13A3 variants may suggest other modifiers are required to promote disease progression. Interestingly, IFN-β therapy induced PAH in a multiple sclerosis patient with a nonsense ATP13A3 mutation (Glu514*) and IFN-β withdrawal improved their PAH symptoms ^30^.

Combined, our results convincingly show that the disease associated ATP13A3 variants present loss-of-function, which based on our knockdown studies would have a major impact on polyamine uptake and homeostasis in ECs.

Importantly, mice harbouring a heterozygous *Atp13a3*^P452Lfs^ variant associated with human PAH developed increased RVSP and right ventricular hypertrophy without altered systemic blood pressure. Also, lung vascular morphometry revealed a significantly higher percentage of fully-muscularized small pulmonary arteries, suggesting that genetic deficiency of *Atp13a3* leads to the development of PAH by affecting small pulmonary vessels. Although the increase in pressure was mild compared to induced PAH models in heterozygous *Atp13a3*^P452Lfs^ variant mice, the PAH phenotype arose spontaneously and reproducibly without requiring a stimulus. This observation is not akin to the reduced penetrance observed in genetic mouse models of *Bmpr2* deficiency, which represents the major the cause of PAH in humans ^31-33^.

In a broader disease context, polyamine dysregulation has been implicated in non-genetic rodent models of pulmonary hypertension (PH) and human PAH. Excessive lung polyamine accumulation was reported in rats with PH induced by either chronic hypoxia ^34, 35^ or monocrotaline (MCT) ^36-38^. However, the different rodent models appear to be associated with different mechanisms of polyamine accumulation. In MCT rats, increased activity of the synthetic enzymes, ODC and AMD, suggests higher polyamine biosynthesis rates ^36, 37^. Administration of the irreversible ODC inhibitor, DFMO, attenuated the increase in mean pulmonary arterial pressure (mPAP) ^38^. Conversely, in hypoxic PH, ^14^C-spermidine accumulation was elevated in lung tissues ^39^, suggesting rates of uptake were increased. Altered polyamine metabolism has also been documented in PAH patients. Metabolomic analyses have reported increased ornithine and putrescine in lung tissues from PAH patients ^40^ and elevated plasma 4-acetamidobutanoate and N-acetyl-putrescine were reported in I/HPAH patients ^41^. Recently, ODC mRNA expression was shown to negatively correlate with mPAP in PAH patients ^42^. Although abnormal polyamine levels are observed in PAH, it is unclear whether this represents changes in the intracellular or extracellular environment and whether various cell types may exhibit different responses. At this juncture, the mechanisms linking polyamine dysregulation and the pathobiology of PAH have not been clarified.

In conclusion, we have demonstrated that ATP13A3 functions as a polyamine transporter and has a functional role in endothelial homeostasis. PAH associated variants exhibited impaired ATP13A3-mediated polyamine transport, contributing to disease-associated cellular phenotypes. These findings shed light for a pathogenic mechanism of ATP13A3 genetic defects leading to loss of function in PAH and provide new insight into a potential role for polyamine dysregulation in the pathobiology of PAH.

## Acknowledgements

We thank S. van Veen and N.N. Hamouda (KU Leuven) for helpful scientific discussions, and the NIHR Imperial Clinical Research Facility for coordinating BOECs preparation. We thank Shruthi Hemanna and Nina Weinzierl (Heidelberg University) for assistance with heart histology. ‘This research was supported by the NIHR Cambridge Biomedical Research Centre (BRC-1215-20014). We wish to thank the BRC for their advice and support in confocal imaging. The views expressed are those of the author(s) and not necessarily those of the NIHR or the Department of Health and Social Care.

## Author Contributions

B.L., M.A., E.L. and J.A.W. conducted experiments. B.L., M.A., E.L., S.M., P.V., P.D.U. and N.W.M. designed experiments and analysed the data. C.V.D.H., V.B. provided lentiviral viral particles. J.W., L.H., M.R.W. provided cell samples. B.L, M.A., E.L., P.V., P.D.U. and N.W.M. drafted the manuscript, and all authors edited the manuscript.

## Data availability

Data are available from the corresponding author by request.

## Supplementary Methods

### Morphometric analysis of the pulmonary vasculature

Pulmonary arteriolar muscularization was assessed on sections of fixed mouse lung tissue (3.5 μm thick) labelled with monoclonal mouse anti–smooth muscle α-actin (αSMA) (clone 1A4; Dako) antibody, followed by polyclonal goat anti-mouse horseradish peroxidase. To detect staining, the ARK kit (Dako) was used in accordance with the manufacturer’s instructions. Antibody staining was visualized using 3-3 diaminobenzidine hydrochloride as substrate-chromogen and counterstained with Carrazzi’s hematoxylin. Pulmonary arteriolar muscularization was assessed by identifying alveolar ducts and categorizing the accompanying intra-acinar artery as non-muscularized, partially muscularized, or fully muscularized by the degree of SMA immunostaining. A minimum of 20 vessels with diameters ranging from 25 to 75 μm were categorized per animal. Invasive hemodynamic measurements as well as morphometric analyses were performed on randomly picked mice and the experimenter was blinded regarding the genotype.

### Wheat germ agglutinin (WGA) staining of heart tissues

Formalin-fixed paraffin-embedded tissue sections (3µm thick) were deparaffinised and rehydrated. The sections were permeabilised with 0.3% Triton in PBS for 20 minutes at RT, blocked with 3% BSA in PBS for another 20 minutes. The tissue sections were then incubated with WGA (1:50, W21404, Invitrogen) in 3%BSA in PBS for 1 hour at RT in the dark. The slides were then mounted with Vectashield HardSetTM Antifade Mounting Medium with DAPI (Vector Laboratories, USA) and visualised using a DMi8 fluorescence microscope (Leica, Germany).

### Transient transfection with siRNA

Cells were maintained in Opti-MEM-I reduced serum media (Invitrogen) for 2 h prior to the addition of DharmaFect1™ (Dharmacon, GE) transfection reagent (4 µl/well in a 6-well plate) with or without siRNA for *ATP13A3* (SASI_Hs02_00356805) or ON-TARGETplus non-targeting Control Pool (siCP) (GE Dharmacon) at a final concentration of 10 nM. The cells were incubated with the siRNA/DharmaFECT1™ for 4 h at 37℃ before the transfection media were replaced with full growth media. Cells were kept in growth media for 24 h before further treatment. Knockdown efficiency was confirmed by assessing mRNA expression RT-qPCR or immunoblotting.

### Transient plasmid DNA transfection

Prior to transfection, HMEC-1 cells were seeded at a density of 250,000 cells/well into a 6-well plate and allowed to adhere overnight. Cells were then incubated in Opti-MEM-I reduced serum media (Invitrogen) for 3 h prior to transfection with 1 µg of pcDNA6.2 expression plasmids encoding either wild type or PAH-associated variant (L675V, M850I, V855M, R858H, L956P) *ATP13A3* with an N-terminal GFP tag. Plasmids were transfected using 9 μl /reaction Lipofectamine LTX with 2.5 μl /reaction Plus reagent (Thermo Fisher Scientific). Cells were incubated with lipoplexes at 37°C for 4 h before being returned to HMEC-1 full growth media. Twenty-four hours post-transfection, cells were trypsinised and reseeded into collagen-coated 4-chambered Nunc™ Lab-Tek™ II Chamber Slides™ (Thermo Fisher Scientific) for immunostaining or other experimental purposes.

### Lentiviral transduction

Lentiviral vectors encoding wild type human *ATP13A3*, the transport dead D498N mutant (D498N) or PAH-associated variants (L675V, M850I, V855M, R858H, L956P) were generated by triple transduction of a transfer plasmid (pCHMWS-ires-puro), an 8.91 packaging plasmid, and a VSV-G envelope plasmid as previously described. ^1^ Prior to transduction, hPAECs were seeded into 6-well plates at a density of 200,000 cells/well and allowed for attaching overnight. The following day, cells were transduced with the lentiviral vectors diluted in EGM-2 supplemented with 2% FBS. hPAECs were transduced with lentiviral particles for 72 h for the optimal transduction. Cells were then used for the following functional assays or lysed directly for extracting protein or RNA.

For stable over-expression of the un-tagged PAH-associated variants, wild type human ATP13A3 or the D498N mutant via lentiviral transduction in HMEC-1 cells, different vector titres were used. Expression was confirmed using immunoblotting and only cells with comparable expression were used. Transduced cells were kept under puromycin (Sigma) selection at the final concentration of 2 µg/mL.

### Stable knockdown of *ATP13A3*

Stable *ATP13A3* knockdown HMEC-1 cell lines were generated using microRNA (miR) based short-hairpin lentiviral vector transduction. Lentiviral particles were produced as described previously ^1^. For the viral transduction, HMEC-1 cells were seeded in a 24-well plate at a density of 100,000 cells/well after which they were incubated with the viral vectors for up to 72 h. Knock-down viral vectors directed at four different target sequences were validated after which the three most potent sequences were selected for further experiments: miR2: AATCACAACAGATTCGTTATTT; miR3: TCAATCGTAAGCTCACTATATT; miR4: AGACCACCTTCGGGTCTTATAT with miR-Fluc: ACGCTGAGTACTTCGAAATGTC used as a negative control. Following transduction, the cells were subjected to selection using Blasticidin (InvivoGen, ant-bl-1) at a concentration of 5 µg/mL.

### Genotyping of the *ATP13A3*^LK726X^ blood outgrowth endothelial cells

*ATP13A3*^LK726X^ BOECs and control BOECs (C4, C7, C35) were grown in 6-well plates at a density of 200,000 cells per well overnight. Genomic DNA (gDNA) of the cells was extracted using the DNeasy Blood & Tissue Kit (Qiagen, West Sussex, UK) in accordance with the manufacturer’s instructions. The resultant gDNA was further PCR amplified with primers (FORWARD: 5’-TGGTTCTTGTGTCACATTTTCAGG-3’; REVERSE: 5’-ACACTCCATTTGCTTCTGTGT-3’) using the AccuPrimeTM Pfx DNA Polymerase kit. PCR products were purified with the Invisorb® Fragment CleanUp kit (Stratec Molecular, Germany) before sanger sequenced (GENEWIZ, UK).

### Proliferation Assay

Cells were seeded in 24-well plates at a density of 30,000/well and left to adhere overnight. Transfection of si*ATP13A3,* siCP or DharmaFECT1™ reagent alone was then performed and cells returned to full growth media afterwards. For assessment of hPAEC proliferation, cells were quiesced in EBM-2/0.1% FBS for 8 h before culturing in EBM-2 media containing 2% FBS (v/v) for six days. Treatments were replenished every 48 h. On day 6, cells were trypsinised with 150 μl/well 0.5% trypsin (Sigma-Aldrich) and quenched with 60 μl/well of the relevant growth media, followed by 90 μl/well of trypan blue (0.4%, Sigma-Aldrich). All 300 μl of cell suspension was transferred into a newly labelled Eppendorf tube, and cells were counted using a haemocytometer. To establish if the level of knockdown was retained, RNA was collected from hPAECs on day 0 (48 h post-transfection) and day 6 followed by assessing *ATP13A3* mRNA expression by qPCR.

### Apoptosis assay

To assess the caspase-3.7 activities, cells were seeded at a density of 150,000/well into 6-well plates and transfected with si*ATP13A3* (Sigma-Aldrich), siCP (GE Dharmacon) or DharmaFECT1™ (GE Dharmacon) alone. For each condition, cells were trypsinised from 6-well plates, reseeded in triplicates into a 96-well plate at a density of 15,000-20,000/well and left to adhere overnight. Cells were quiesced in EBM-2/0.1% FBS for 24 h before culturing in EBM-2/0.1%FBS for 16 h. For measuring caspase-3/7 activities, 100 μl Caspase-Glo® 3/7 Reagent (G8091 Promega) was added into each well, then incubated and mixed on a plate shaker in the dark for 15 min at room temperature. The entire 200 μl from each well was transferred into the corresponding well of a white-walled 96-well plate and luminescence was read in a GloMax® luminometer (Promega).

### Endothelial permeability assay

This assay measures the transit of horseradish peroxidase (HRP) across endothelial monolayers seeded in transwell inserts. hPAECs subjected to siRNA transfection were trypsinised and reseeded at a density of 50,000 cells/insert into Corning® Transwell® chambers with polyester membrane cell culture inserts (Corning) and allowed to attach for 30 min before adding 1ml of EGM2 supplemented with 2% FBS (Promocell) to the lower 24-well plate chamber. hPAECs were incubated overnight and then serum-starved for 6 h by replacing the media in the inserts with 200 μl, and the bottom chambers with 1ml of EBM2/0.1% FBS. Following serum-starvation, media were then replaced with 2% FBS supplemented EBM2 with or without the addition of 1 U/ml Thrombin (Sigma) and incubated for 1 h at 37°C. In the meantime, 0.05 M Phosphate Citrate Buffer was prepared by dissolving one capsule of Phosphate-Citrate buffer with Sodium Perborate (Sigma) into 100 ml of PBS. The o-Phenylenediamine dihydrochloride (OPD) developing solution was prepared by adding one OPD (Sigma) tablet into 50 ml Phosphate-Citrate buffer. Prior to the assessment of permeability, media in the inserts was replaced with 100 μl of EBM2 supplemented with 2% FBS containing 25 nM HRP(Sigma) with or without the addition of 1 U/ml Thrombin. 3 x 15 μl medium from the lower chambers were collected into a new 96-well plate at the following time points: 0min, 15min, 30min, 1 h, 1.5 h and 2 h. 150 μl/well of OPD buffer was then added into the 96-well plate and the absorbance measured at 490 nm over 10 to 20 min.

### Cytotoxicity Assay

Cells were seeded into 96 clear-well F-bottom plates at a cell density of 2.5 X 10^4^ cells per well and incubated overnight (5% CO2 and at 37°C) to attach to the bottom of the wells. Doses of the screened compounds were made in DMEM cell culture medium and 100 μL of the dilutions were added to the corresponding wells. After an overnight incubation, the cells were washed twice with PBS. Finally, 50 μL of 4-Methylumbelliferyl heptanoate (MUH reagent) (Sigma), dissolved in PBS to a final concentration of 100 μg/mL, was added per well. The cells were then incubated at 37°C in the dark for 45 min. The fluorescence was measured using a multi-mode plate reader (Flexstation3, Molecular Devices) with excitation at 355 nm, emission at 460 nm and cut-off at 455 nm. Results were normalised to the untreated control.

### RNA extraction and quantitative reverse transcription-PCR

Total RNA was extracted using RNeasy Mini Kit buffers (Qiagen, West Sussex, UK) and Silica Membrane Mini Spin Columns (EconoSpin) following the manufacturer’s instructions. Equal amounts of RNA (∼1 μg) were then reverse transcribed into cDNA using a High Capacity Reverse Transcriptase kit (Applied Biosystems). 2 μl cDNA, 1.8 μl associated premixed primer sets (final concentration in mix = 200nM), 5 μl 2X SYBR Green JumpStart Taq ReadyMix (Sigma-Aldrich), 0.2 μl ROX reference dye (Invitrogen) and 1 μl DEPC-treated water were prepared into one well of a MicroAmp® Optical 384-Well Reaction Plate (Applied Biosystems) before loading on a QuantStudio 6 Flex Real-Time PCR System (Applied Biosystems). Amplification reactions were initiated with a 2-min pre-incubation at 95°C, followed by 50 amplification cycles of 30-second denaturation at 95°C, 30 seconds annealing at 55°C and 30-seconds extension at 72°C. Melt curve analysis was performed to rule out nonspecific amplification, and no-template controls were included. Primers for human genes (supplement table 2) were designed using Primer-BLAST (http://www.ncbi.nlm.nih.gov/tools/primer-blast/), and primer efficiency confirmed before use. The relative expression levels of target genes were calculated using the 2^^-(△△Ct)^ method by normalizing to the stably expressed housekeeping genes, Beta-2-Microglobulin (*B2M*) or β-actin (*ACTB*). Differences in gene expression are presented as the fold change relative to control. Relative expression of target genes in different cell lines or mouse tissues was determined as 2^^-(CTtarget -CThousekeeping)^.

### Cellular polyamine measurement

### Extraction of aqueous metabolites

Following siRNA transfection or lentiviral transduction, cells were washed twice with phosphate-free buffer (162 mM ammonium acetate 7.4) before being lysed with 4:1 methanol: water lysis buffer. The resulting cell lysate was sonicated for 5 min in a water bath sonicator followed by an additional 5-min sonication if obvious cell debris was still present. The residual debris was then removed by pelleting the cell homogenate at 21000 x g for 10 min at RT, and the supernatant was carefully transferred into new tubes.

### LC-MS sample preparation for analysis of polyamines

Aqueous extracts of cells were dried using a centrifugal evaporator (Savant, ThermoFisher) and reconstituted in 50 µl of 10 mM ammonium acetate containing 2 µM [^13^C_10_, ^15^N_5_] adenosine monophosphate, 10 µM succinic acid [^13^C10], 10 µM d8 putrescine and a 1 in 5000 diluted [U^13^C_10_, U^15^N_5_] mixture of amino acids (all purchased from Sigma Aldrich with the exception of the d8 putrescine which was obtained from CDN isotopes) as internal standards. The resulting solution was vortexed then sonicated for 5 min. followed by brief pulsed centrifugation to recover the maximum volume. After centrifugation the supernatant was transferred with an automatic pipette into a 300 µl glass vial (Chromacol) and capped (Agilent) ready for analysis.

### LC-MS analysis of polyamines

For all analysis a Q Exactive Plus orbitrap coupled to a Vanquish Horizon ultra-highperformance liquid chromatography system was used. LC analysis was carried out using an ACE Excel C18-PFP column (150 × 2.1 mm, 2.0 µm, Hichrom). Mobile phase A consisted of water with 0.1% formic acid with 10 mM ammonium formate and mobile phase B was acetonitrile with 0.1% formic acid. For gradient elution mobile phase B was held at 0% for 1.6 min followed by a linear gradient to 30% B over 4.0 min, a further increase to 90% over 1 min and a hold at 90% B for 1 min with re-equilibration for 1.5 min giving a total run time of 6.5 min. The flow rate was 0.5 mL/min and the injection volume was 2 µL. The needle wash used was 1:1 water:acetonitrile.

Samples were run using electrospray ionisation in positive ion mode only. Source parameters used for the orbitrap were a vaporizer temperature of 450°C and ion transfer tube temperature of 320°C, an ion spray voltage of 3.5 kV and a sheath gas, auxiliary gas and sweep gas of 55, 15 and 3 arbitrary units respectively with an S-lens RF (radio frequency) of 50%. A full scan of 60-900 m/z was used in positive ion mode at a resolution of 70,000 ppm.

### LC-MS data processing

Data were acquired, processed and integrated using Xcalibur (Version 4.1, Thermo Fisher Scientific). Retention times and accurate masses for putrescine, acetyl putrescine, spermine and spermidine were validated against injections of known external standards (obtained from Sigma Aldrich). Peak areas corresponding to metabolite levels were manually quantified and normalised to the appropriate internal standard and presented as relative areas.

### Immunoblotting

Cells were snap-frozen and lysed in SDS-lysis buffer (125mM Tris (pH 7.4), 2% SDS, 10% glycerol) containing an EDTA-free protease inhibitor cocktail (Roche) or by using RIPA buffer (Thermo Fisher) containing protease inhibitors (Sigma). Protein concentrations were assessed using a modified Lowry assay (Bio-Rad DC Assay, Bio-Rad) or Pierce BCA Protein Assay Kit (Thermo Fisher). Primary antibodies for ATP13A3 (amino acids 488-631, Sigma, HPA029471). and GAPDH (Sigma, G8795) were dissolved at a dilution of 1:1000 and 1:5000 respectively in 1% BSA/TBS-tween and incubated overnight at 4°C. The following secondary antibodies, IgG HRP-linked Anti-rabbit (Cell Signalling, 7074S) and IgG HRP-linked Anti-mouse (Cell Signalling, 7076S), were dissolved in 5% milk/TBS-tween solution at a dilution of 1:5000 for 1 h, Bands were detected using SuperSignal™West Pico PLUS chemiluminescent Substrate (Thermo Fisher) or incubated with enhanced chemiluminescence reagent (GE Bioscience) and visualised by on the Bio-Rad Chemidoc ™ MP imaging system or using X-ray film (GE healthcare)..

### Immunofluorescence staining

Prior to cell seeding, 4-chambered Nunc™ Lab-Tek™ II Chamber Slides™ (Thermo Fisher Scientific) were pre-coated with 500μl/chamber Type I Rat Tail Collagen (BD Biosciences) for 90 min before washing three times with PBS. Following siRNA or plasmid DNA transfection, cells were trypsinised and reseeded into collagen-coated 4-chambered slides and left to adhere overnight. Cells were washed with PBS and fixed in 4% (v/v) Paraformaldehyde (Sigma) at RT for 10 min, then permeabilised with 0.05%(v/v) Saponin in 0.5% BSA/PBS for 20 min and blocked in 0.5% BSA/PBS for 1 h. The chambers were removed and the slides incubated overnight with the primary antibodies listed below (supplement table 3). Cells were then washed three times with PBS and incubated with the corresponding fluorescent antibodies (Online Table III) at RT for 1 h. Cells were then washed twice with PBS before mounting with Vectamount™ medium containing DAPI (Vector Laboratories) and imaging on a Leica Sp5 confocal microscope platform (Leica microsystem).

### Confocal Microscopy of Cellular BODIPY-PUT Uptake

Cells were seeded in a 12-well plate at a density of 10,000 cells/well on coverslips. The following day, they were then then treated with the polyamine-BODIPY probes as described earlier. The medium was then aspirated, the cells washed with PBS and fixed with 4% paraformaldehyde for 30 min at 37°C after which the cells were once again washed with PBS. Next the cells were permeabilized with 0.1% Triton-X and blocked for 1 h with 0.1 M glycine and then for 1 h in PBS containing 0.1% Tween (PBS-T), 10% FBS and 1% BSA. Next the cells were stained with DAPI for 15 min, the slides were washed with PBS-T and glued onto slides using the Alexa FluorSave reagent and left to dry. Slides were imaged using the LSM780 confocal microscope.

## Supplementary figures and tables

**Supplement Figure 1.**
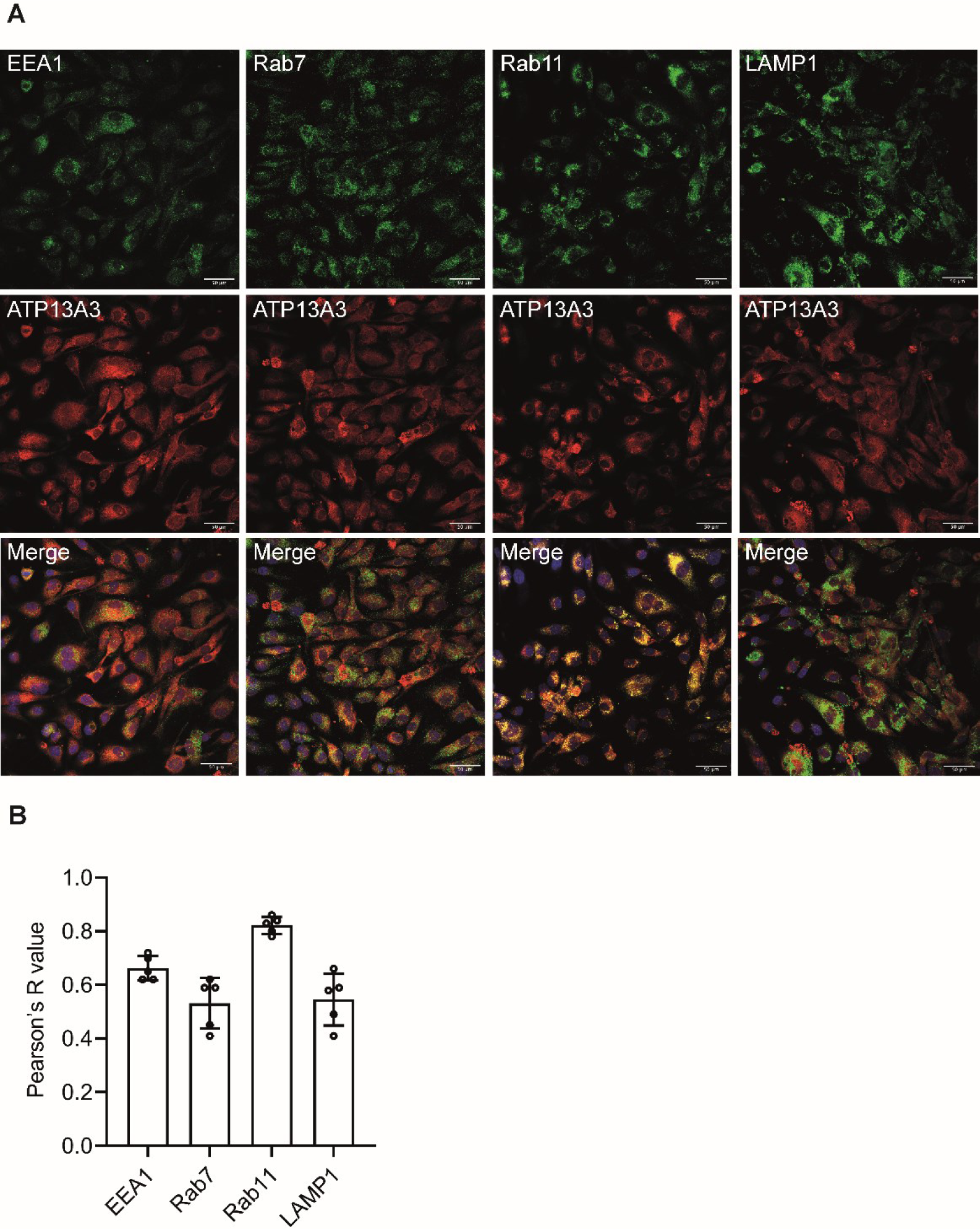
Analysis of endogenous ATP13A3 cellular compartmentalisation in HMEC-1 cells. (A) Confocal images (40X, scale bar = 10 µm) of HMEC-1 co-stained with anti-ATP13A3 and antibodies against either EEA1, Rab7, Rab11 or LAMP1. (B) Pearson’s coefficients of ATP13A3 to different endosomal markers in HMEC-1.

**Supplement Figure 2.**
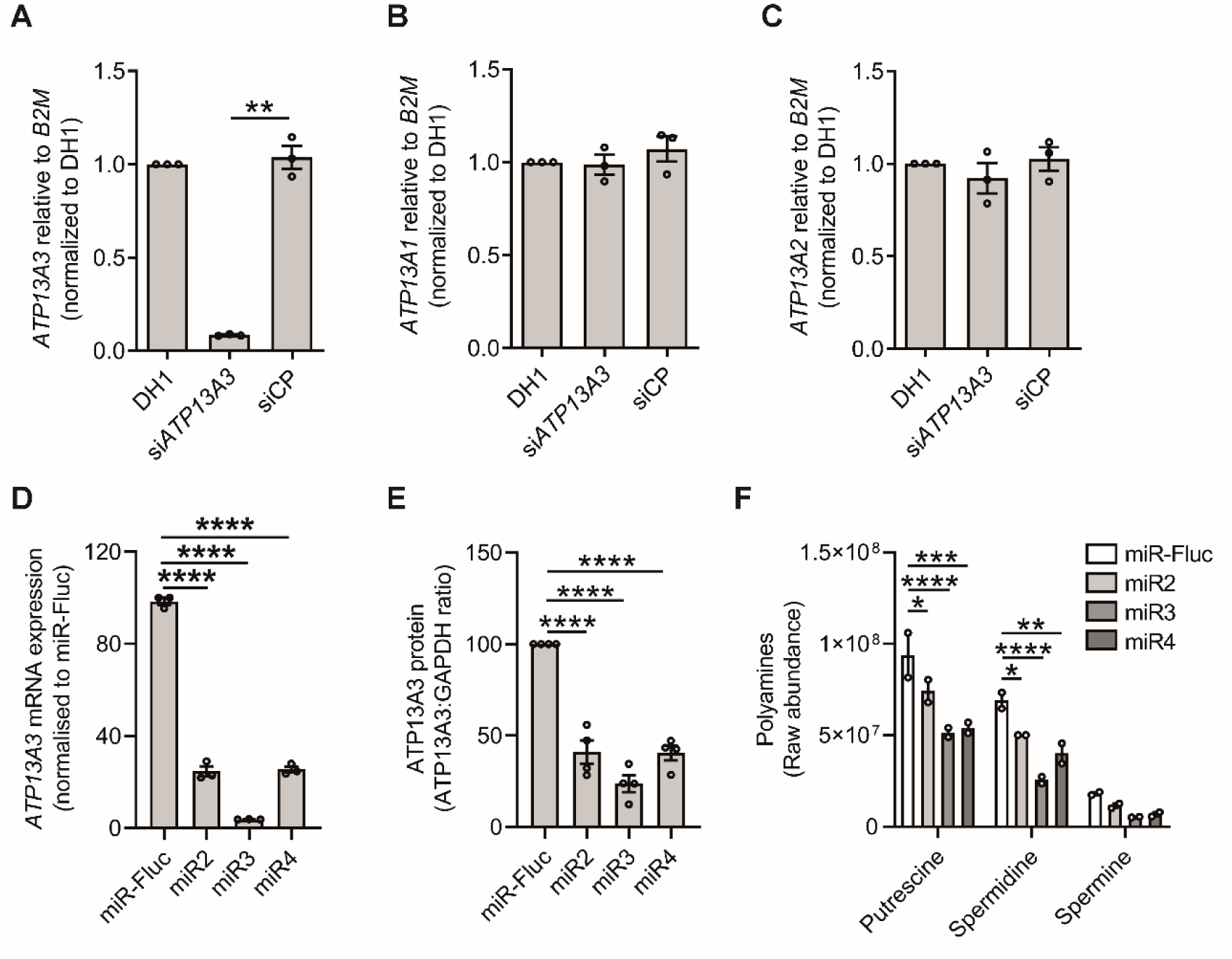
mRNA expression of P5-type ATPases in *ATP13A3* silenced hPAECs and HMEC-1 cells. Expression of (A) *ATP13A3* (B) *ATP13A1* (C) *ATP13A2* mRNAs in hPAECs transfected with DharmaFECT1™ (DH1) alone, si*ATP13A3* or non-targeting siRNA control (siCP) (n=3 experiments). (D,E) *ATP13A3* (D) mRNA and (E) protein expression in HMEC-1 cells stably expressing miR2-4 and miR-Fluc. Data (n=3 experiments) are mean ± SEM expressed relative to (A-C) DH1 or (D,E) miR-Fluc. (F) Metabolomic analysis of the raw abundance of basal intracellular putrescine, spermidine and spermine levels in HMEC-1 cells stably expressing miR2-4 and miR-Fluc (n=2 experiments). Data were analysed using a One-way ANOVA with Tukey’s *post hoc* test for multiple comparisons. **P<0.05, ***P<0.01, ****P<0.0001.

**Supplement Figure 3.**
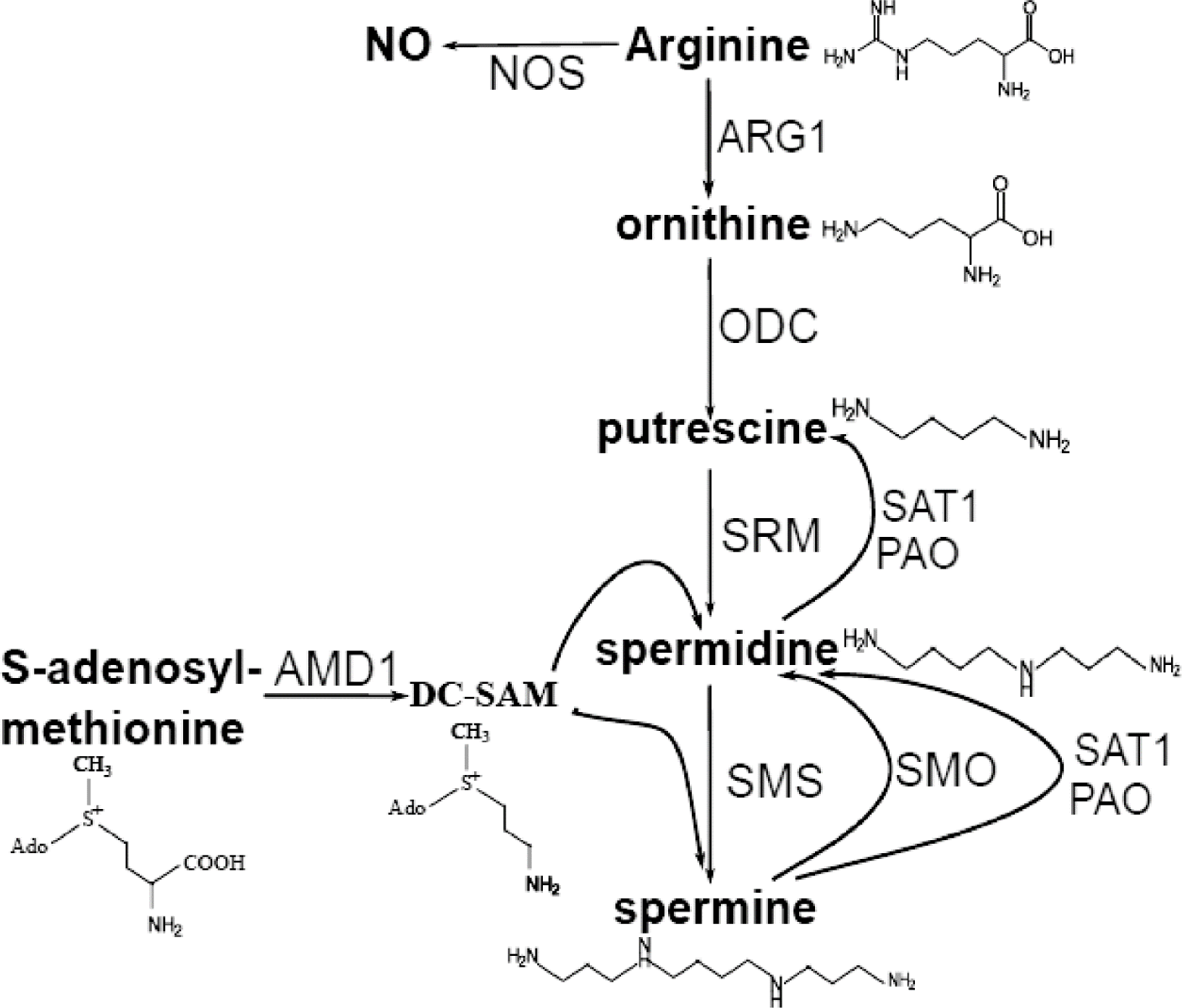
Schematic of cellular polyamine metabolism pathways. Arginase 1 (ARG1) mediates the conversion of arginine to ornithine, which is then converted to putrescine by ornithine decarboxylase (ODC). Putrescine is then converted to spermidine by spermidine synthase (SRM), which can then be converted to spermine by spermidine synthase (SMS). Conversely, spermine is converted to spermidine by spermine oxidase (SMO/SMOX), spermine/spermidine acetyltransferase (SAT1) and polyamine oxidase (PAO), the latter two also mediating conversion of spermidine to putrescine. In addition, S-adenosylmethionine (AMD1) activity can generate spermidine or spermidine via decarboxylaton of S-adenosyl-methionine to the intermediate, methyl-S-adenosylthiopropylamine (DC-SAM), which can be converted to spermidine by SRM.

**Supplement Figure 4.**
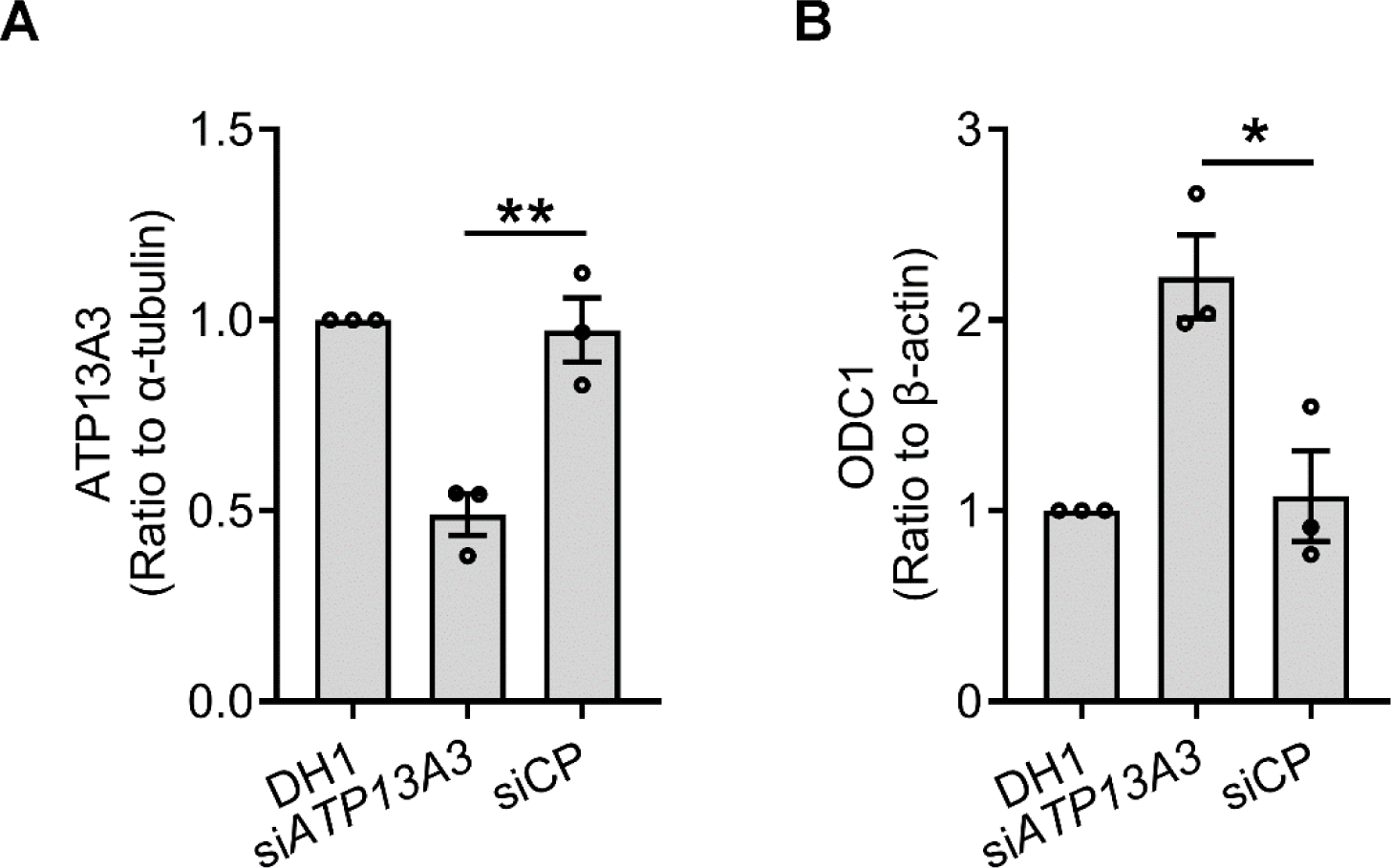
Densitometric analysis of (A) ATP13A3 and (B) ODC normalised relative to α-tubulin and β-actin respectively and as fold-change relative to DH1. (Blots are shown in Figure 3A)

**Supplement Figure 5.**
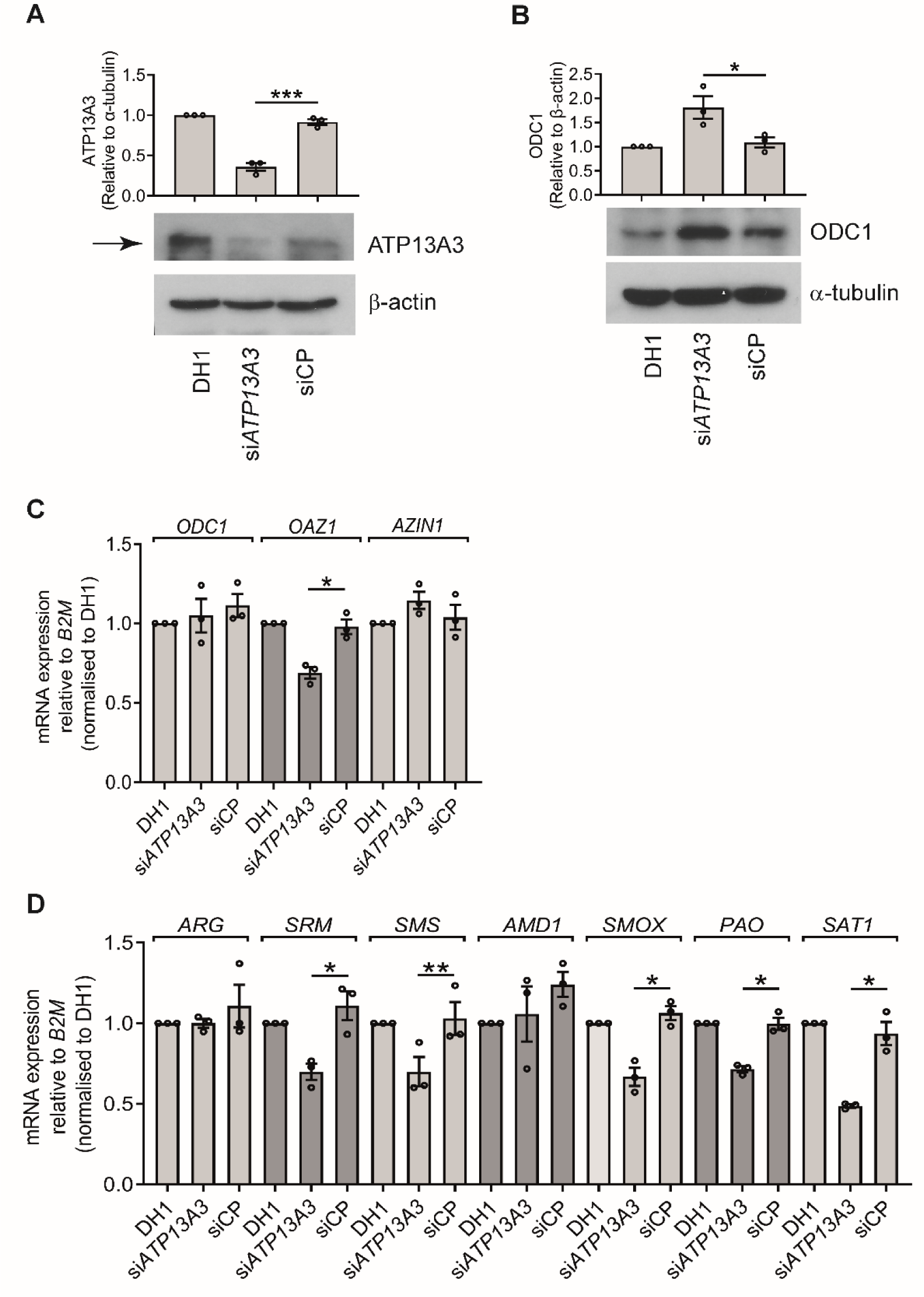
*ATP13A3* silencing disrupted polyamine metabolism in BOECs. Immunoblotting of (A) ATP13A3 and (B) ODC1 in control BOECs transfected with DharmaFECT1™ (DH1) alone, si*ATP13A3* or non-targeting siRNA control (siCP). Densitometric analysis of ATP13A3 and ODC relative to α-tubulin and β-actin respectively are shown above the blots. (C,D) Transcriptional alteration of (C) *ODC1, OAZ1* and *AZIN1,* (D) catabolic enzymes (*SMOX*, *PAO*, *SAT1*) mRNA and (E) polyamine biosynthesis enzymes (*ARG1*, *SRM*, *SMS*, *AMD1*) in BOECs transfected with DH1, si*ATP13A3* or siCP. Data (n=3 experiments) are mean ± SEM and are fold-change relative to DH1. Data were analysed using a One-way ANOVA with Tukey’s post hoc test for multiple comparisons. *P<0.05, **P<0.01 and ***P<0.001

**Supplement Figure 6.**
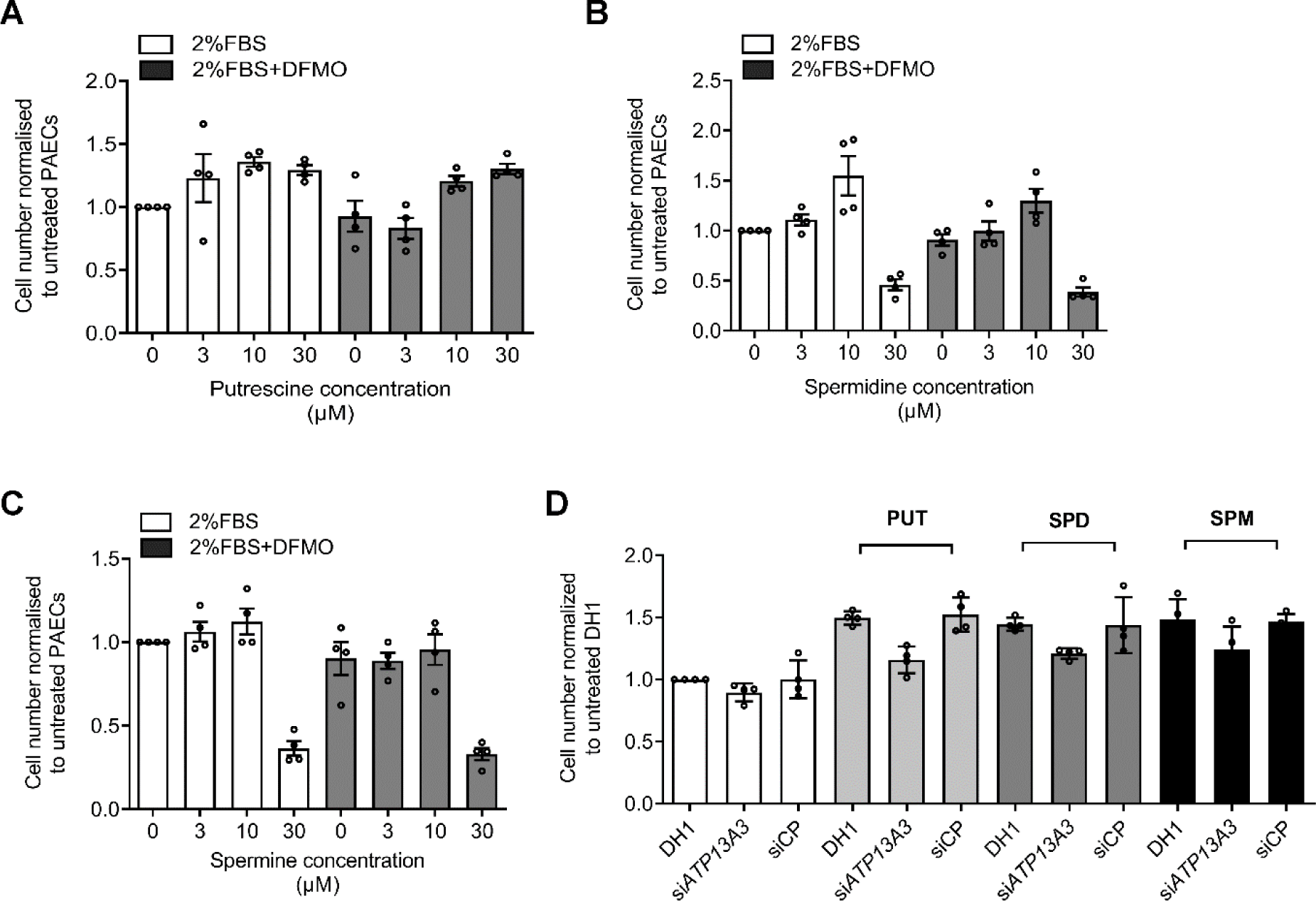
Effects of supplementation with polyamines on PAEC proliferation. Proliferation of hPAECs following 6-day incubation in EBM2 with 2% alone, or supplemented with (A) putrescine, (B) spermidine or (C) spermine, with or without DFMO at the indicated concentration. Media were replenished every other day. (D) PAECs transfected with DharmaFECT1™ (DH1), si*ATP13A3* or siCP were treated with 10µM of either putrescine (PUT), spermidine (SPD) or spermine (SPM) and assessed for proliferation at day 6. Data for (A-C) are presented as fold change relative to untreated control and (D) normalised to untreated DH1. Mean ± SEM are shown and data were analysed by one-way ANOVA with Tukey’s post hoc test for multiple comparisons.

**Supplement Figure 7.**
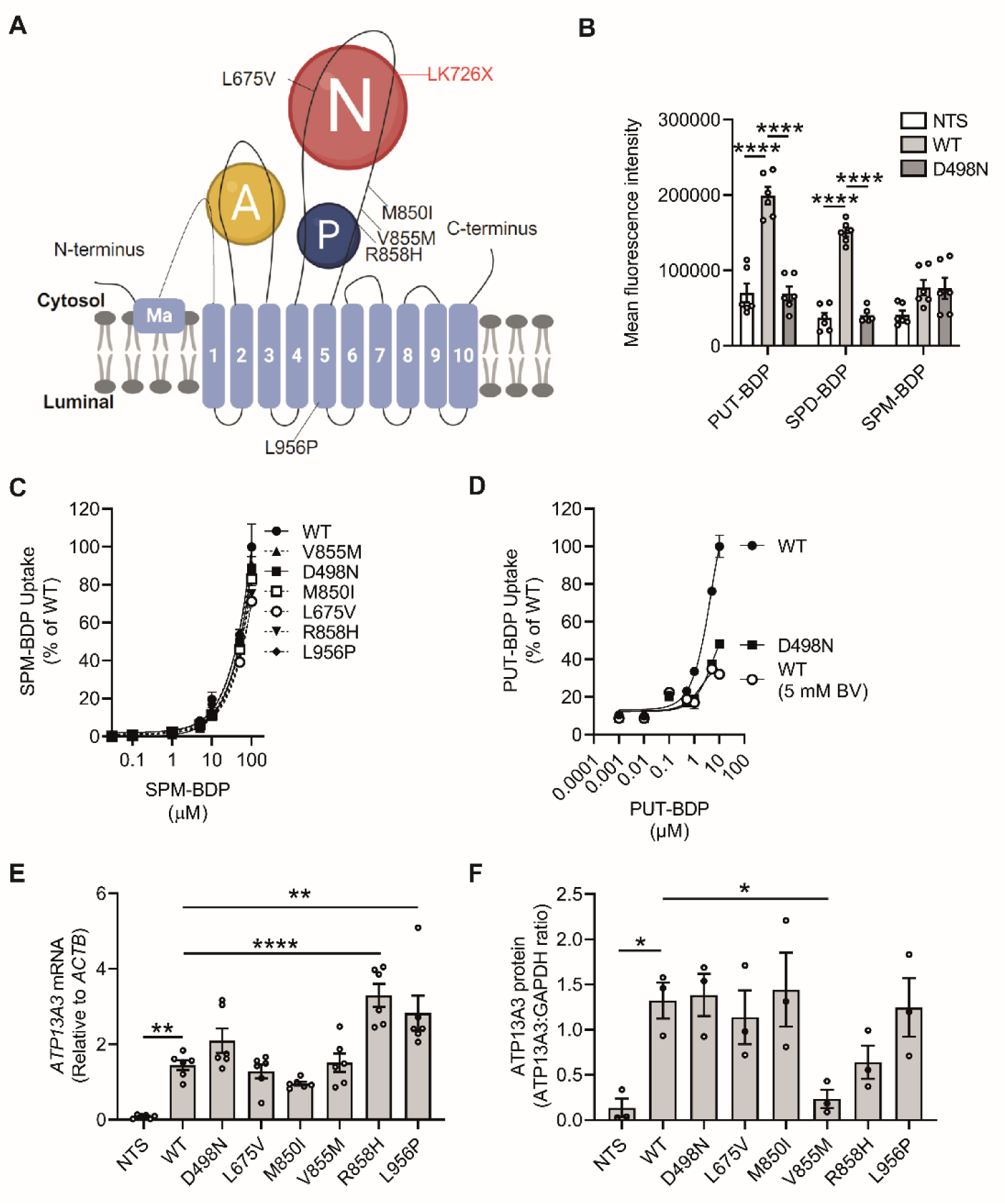
**ATP13A3 expression and validation of WT-ATP13A3 as a polyamine transporter. (**A) Predicted ATP13A3 topology showing the three cytosolic domains, the Actuator domain (A, yellow), nucleotide domain (N, red) and the phosphorylation domain (P, blue). The protein contains 10 membrane-spanning segments, responsible for substrate binding and provision of structural support. An additional N-terminal segment (Ma) is embedded in the membrane. The location of disease related missense (black) and frameshift (red) variants are indicated on this structure. (B) Flow-cytometry analysis (n=6 experiments) of PUT-BDP, SPD-BDP and SPM-BDP uptake after 30 minutes exposure in HMEC-1 cells stably expressing ATP13A3 wild-type (WT) or the D498N mutant compared to non-transduced (NTS) cells. (C) Flow-cytometry analysis for assessment of cellular uptake increasing concentrations of SPM-BDP (n=2 experiments) after 30 minutes exposure. Data are normalised to WT. (D) PUT-BDP uptake by HMEC-1 stably overexpressing ATP13A3 WT and D498N protein and in the WT cells treated with 5 mM Benzyl Viologen, a polyamine transport inhibitor. (E,F) The expression of *ATP13A3* (E) mRNA (n=3 experiments) and (F) protein (n=3 experiments) in non-transduced (NTS) HMEC-1 cells compared to those stably expressing untagged ATP13A3 WT (WT), a transport dead mutant (D498N) or five PAH-associated variants (L675V, M850I, V855M, R858H and L956P). (F) mRNA levels are expressed relative to *ACTB* and (G) protein levels were normalised to GAPDH (see Figure 5A). Mean ± SEM are shown and data were analysed by one-way ANOVA with Tukey’s post hoc test for multiple comparisons. *P<0.05, **P<0.005 and ****P<0.0001

**Supplement Figure 8.**
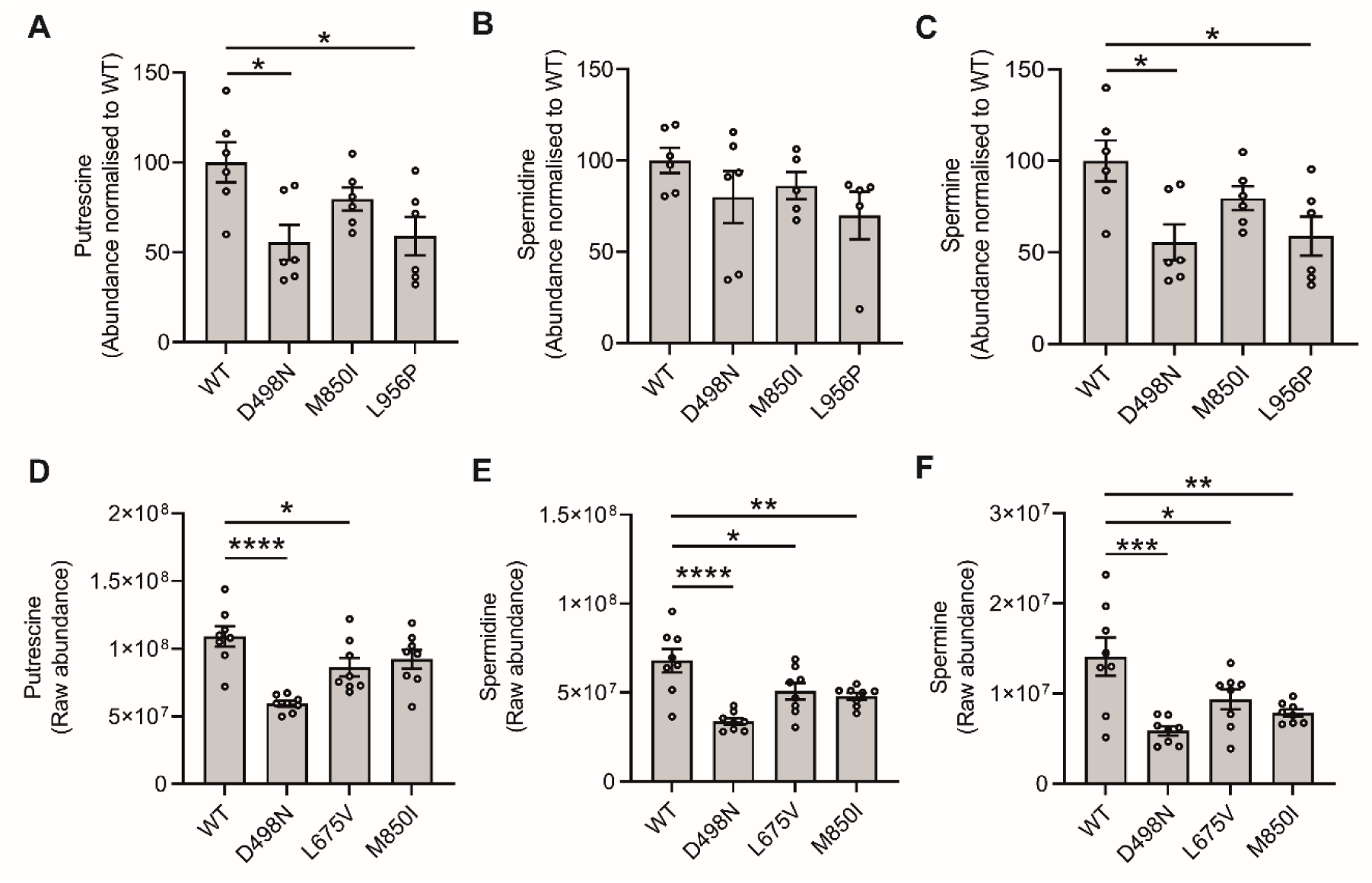
Basal polyamine levels are not reduced in HMEC-1 stables, but are reduced in SH-SY5Y cells overexpressing the M850I ATP13A3 variant. (A-C) Metabolomics data showing the normalised basal intracellular levels of (A) putrescine, (B) spermidine and (C) spermine in HMEC-1 cells over-expressing ATP13A3 WT (WT), D498N and the L675V and M850I PAH missense variants. (D-F) Metabolomics data showing the total basal intracellular levels of (D) putrescine, (E) spermidine, and (F) spermine in SH-SY5Y cells over-expressing ATP13A3 WT, transport dead mutant (D498N) and the L675V and M850I PAH missense variants. Mean ± SEM are shown and data were analysed by one-way ANOVA with Tukey’s post hoc test for multiple comparisons. *P<0.05, **P<0.005, ***P<0.001 and ****P<0.0001.

**Supplement Figure 9.**
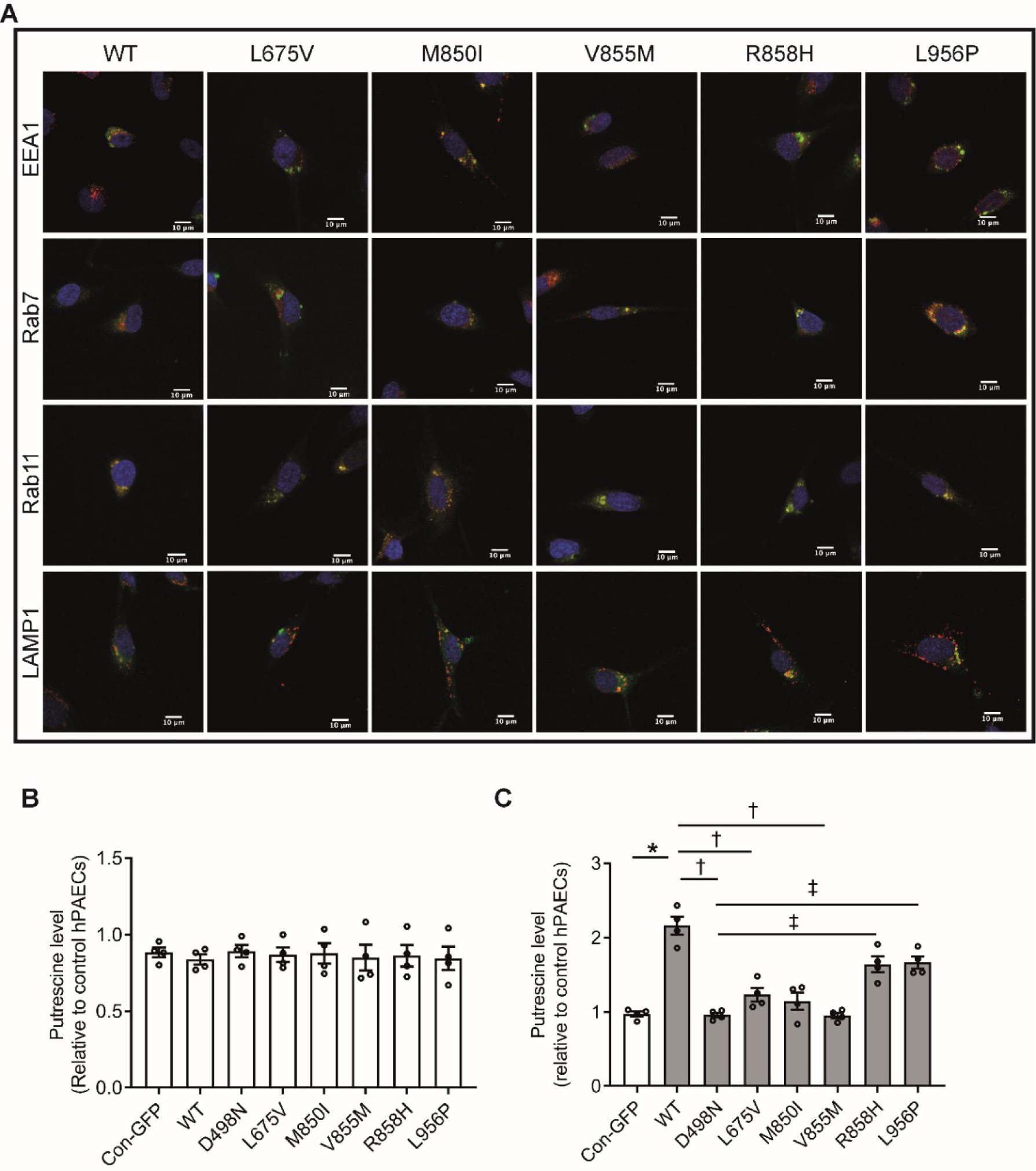
PAH associated mutations impair ATP13A3 mediated putrescine uptake in endothelial cells. (A) Confocal images (63X, scale bar = 10 µm) of HMEC-1 transiently over-expressing GFP-tagged wild type (WT) or PAH-associated variants (L675V, M850I, V855M, R858H, L956V) GFP-*hATP13A3* co-stained with antibodies against either EEA1, Rab7, Rab11 or LAMP1. (B-C) Cellular putrescine in hPAECs overexpressing lentiviruses encoding ATP13A3 WT, D498N, PAH-associated variants or GFP-tagged empty vectors. Cells were incubated overnight in EBM2 containing: (B) 2%FBS alone or (C) with 1mM Putrescine. Data (n=4 experiments) are polyamine peak area ratio relative to sample protein concentration normalised to the GFP-empty vector. Data are mean ± SEM analysed by one-way ANOVA with Tukey’s post hoc test for multiple comparisons *P<0.05 compared with con-GFP. ^†^P<0.05 compared with WT, ^‡^P<0.05 compared with D498N.

**Supplement Figure 10.**
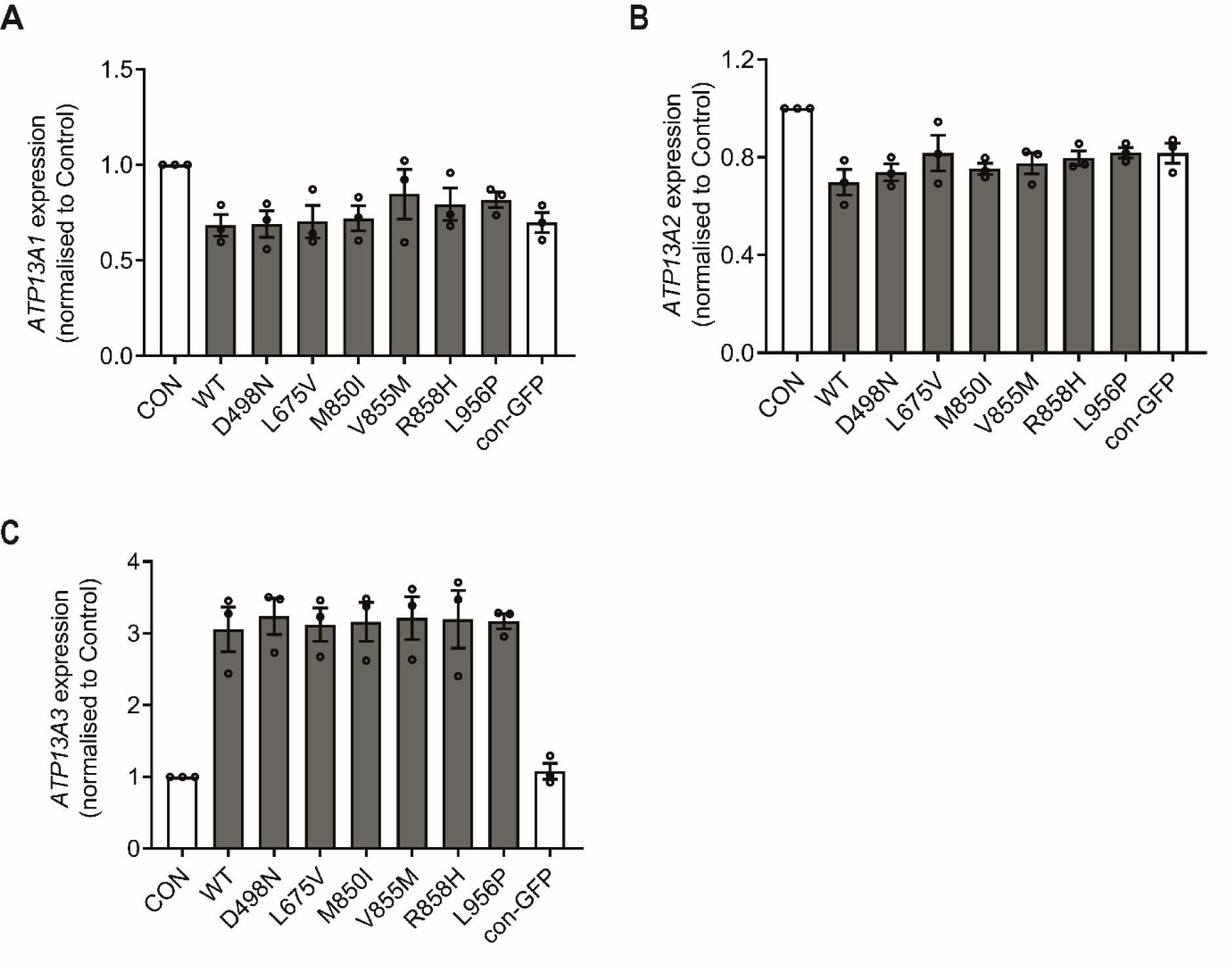
mRNA expression of P5-type ATPases in hPAECs transiently overexpressing WT and PAH-associated ATP13A3 variants. mRNA expression of (A) *ATP13A1* (B) *ATP13A2* (C) *ATP13A3* of control hPAECs (CON), and hPAECs transiently transduced with lentiviruses encoding GFP-tagged ATP13A3 WT (WT), transport dead mutant (D498N), PAH-associated variants (L675V, M850I, V855M, R858H, L956P) or non-targeted control (con-GFP). Data (n=3 experiments) are mean ± SEM and presented as fold-change relative to DH1.

**Supplement Figure 11.**
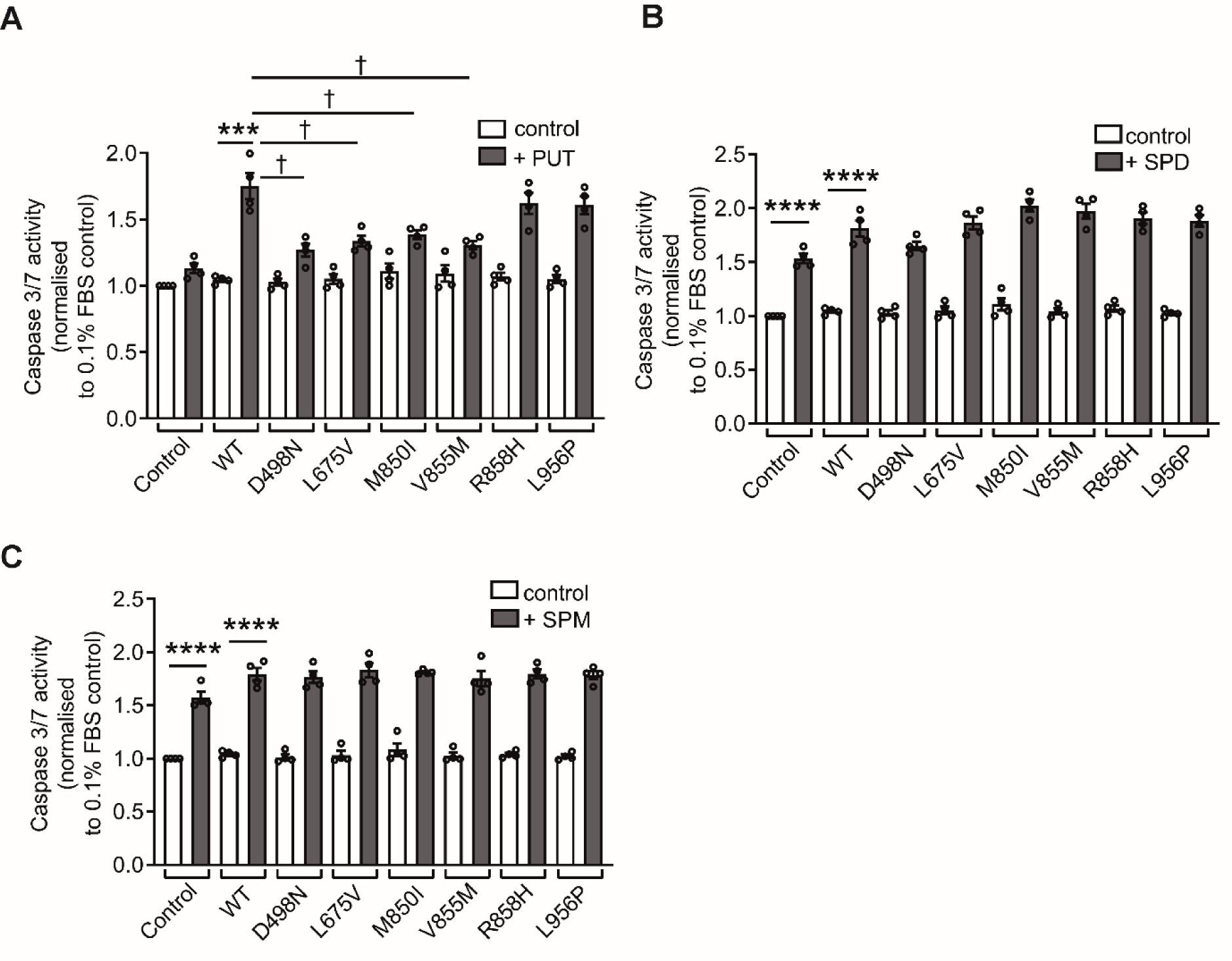
ATP13A3 regulates the endothelial response to polyamine-induced stress. Apoptosis of hPAECs transduced with lentiviruses encoding GFP-tagged *ATP13A3* WT, D498N mutant and PAH-associated variants (L675V, M850I, V855M, R858H, L956P). Cells were cultured overnight in EBM2 containing 0.1%FBS with or without (A)10mM putrescine (PUT), (B)1mM spermidine (SPD) or (C)1mM spermine (SPM) and apoptosis assessed by Caspase-Glo®3/7 assay. Data (n=4 experiments) are mean ± SEM of the fold change relative to hPAECs cultured in 0.1% FBS. ***P<0.001, ****P<0.0001 unpaired t-test compared with cells in 0.1%FBS. ^†^P<0.05 one-way ANOVA with Dunnett’s test for multiple comparisons with WT

**Supplement Figure 12.**
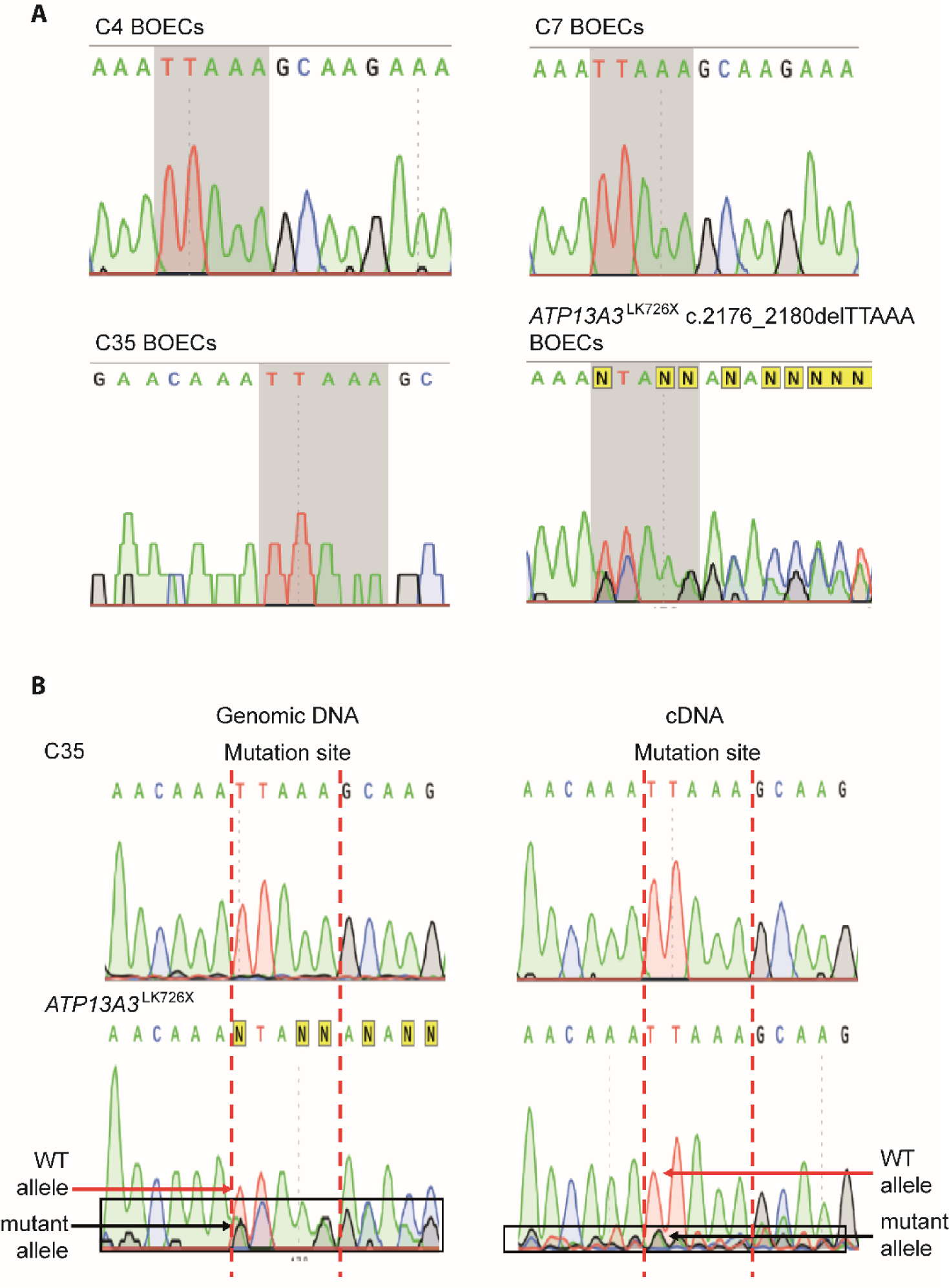
*ATP13A3* LK726X frameshift variant induces nonsense-mediated mRNA decay (NMD) in BOECs. (A) Sanger sequencing data of the PCR products amplified from the genomic DNA of control BOECs (C4, C7, C35) and *ATP13A3*^LK726X^ BOECs. *ATP13A3*^LK726X^ BOECs showed the heterozygous deletion of TTAAA at the c.2176_2180 variant site. (B) Sanger sequencing data of the PCR products amplified from the genomic DNA (gDNA) and cDNA of control BOECs (C35) and *ATP13A3*^LK726X^ BOECs. Wild type (WT) and variant alleles are indicated by red and black arrows respectively. The peak height ratio of the variant allele relative to the wild type (WT) allele at the variant site is lower in the cDNA sequencing chromatogram than in the gDNA sequencing chromatogram, indicating LK726X may induce NMD in the *ATP13A3*^LK726X^ BOECs.

**Supplement Figure 13.**
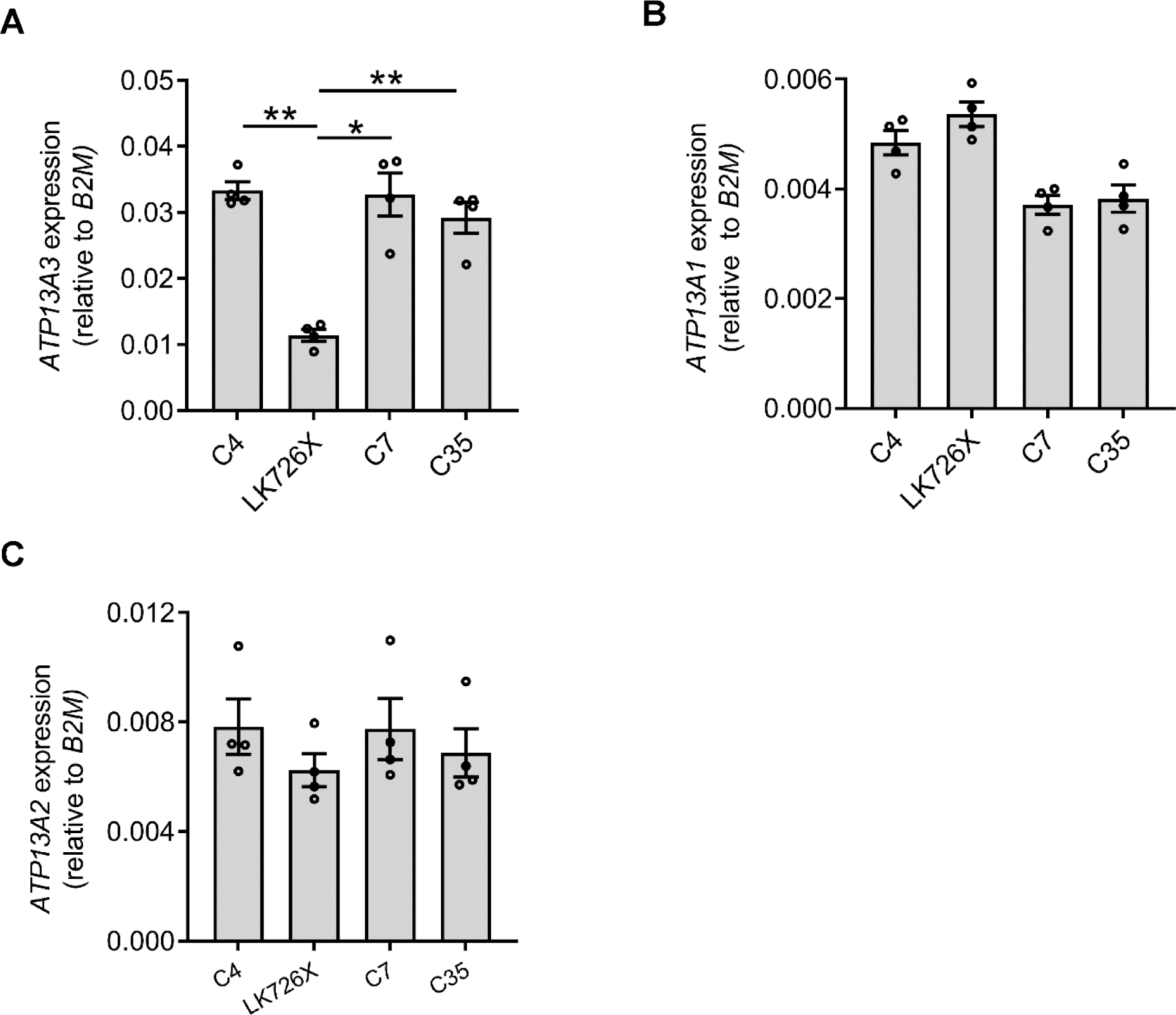
*ATP13A1-3* mRNA expression in control and ATP13A3^LK726X^ BOECs. mRNA expression of (A) *ATP13A3,* (B) *ATP13A1* and (C) *ATP13A2* in control BOECs (C4, C7, C35) and *ATP13A3*^LK726X^ (LK726X) BOECs. Data (n=4 experiments) are mean ± SEM presented as relative expression normalised to *B2M*. Data were analysed using a One-way ANOVA with Tukey’s post hoc test for multiple comparisons. *P<0.05, **P<0.01.

**Supplement Figure 14.**
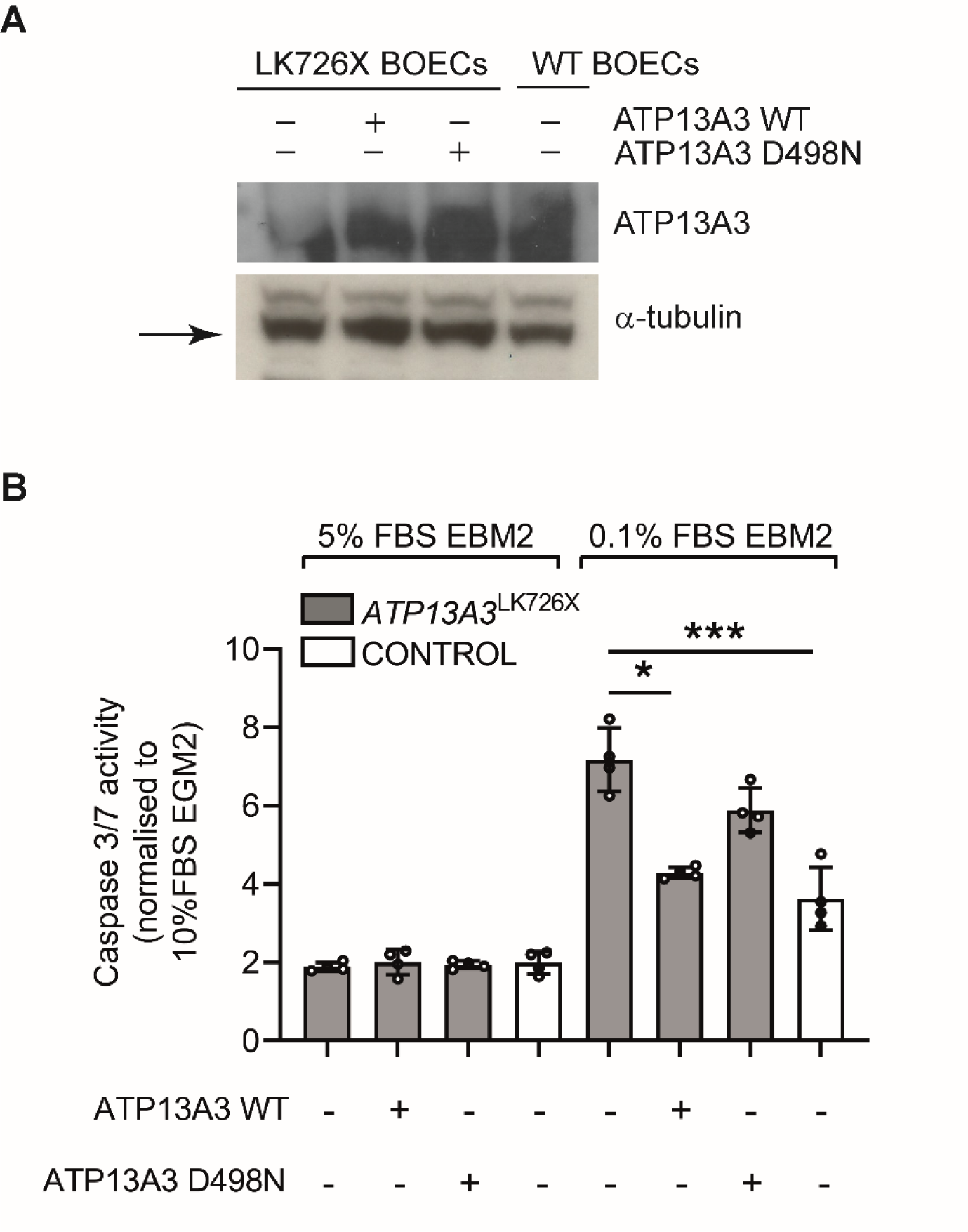
Wild type *ATP13A3* overexpression rebalances the pro-apoptotic phenotype in *ATP13A3*^L726X^ BOECs. *ATP13A3*^LK726X^ BOECs were transiently transduced with or without lentiviruses encoding ATP13A3 WT or the D498N mutant. (A) ATP13A3 protein expression was assessed by immunoblotting. Cells cultured in in EBM2 supplemented with 5%FBS or 0.1%FBS were assessed for apoptosis by Caspase-Glo®3/7 assay (B). Data (n=4 experiments) are mean ± SEM analysed by one-way ANOVA with Tukey’s post hoc test for multiple comparisons *P<0.05 compared with con-GFP. *P<0.05 ***P<0.001 compared with *ATP13A3*^LK726X^ BOECs cultured in 0.1%FBS EBM2.

**Supplement Figure 15.**
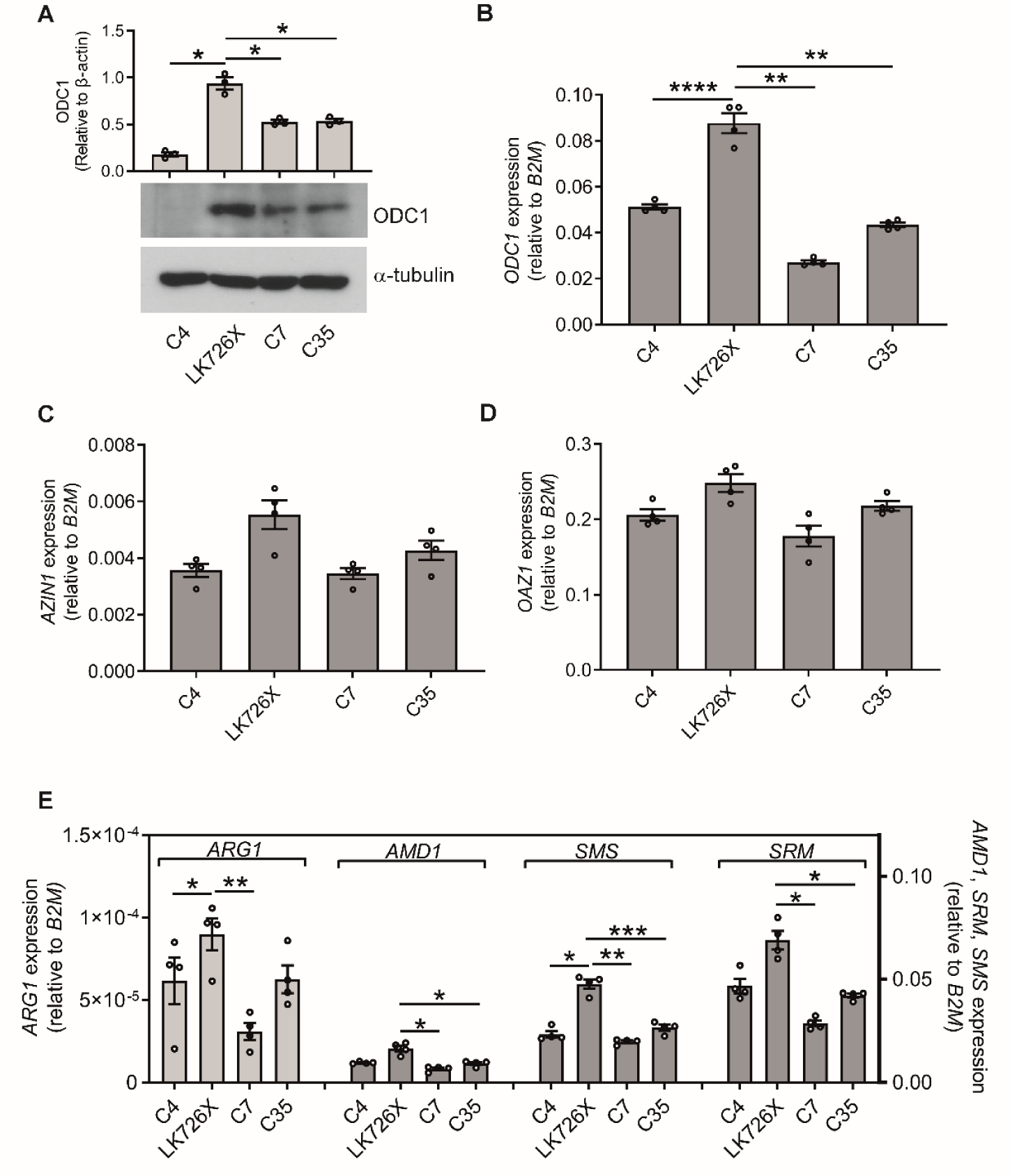
The *ATP13A3* LK726X frameshift mutation alters polyamine metabolism pathways. (A) Immunoblotting of ODC1 in control BOECs (C4, C7, C35) and *ATP13A3*^LK726X^ (LK726X) BOECs. Densitometric analysis of ODC1 and β-actin was performed. (B)*ODC*, (C) *AZIN* and (D) *OAZ1* mRNA of BOECs is presented as expression relative to *B2M*. (E) mRNA expression of polyamine biosynthesis related enzymes (*ARG1, AMD1, SRM, SMS*) presented as expression relative to *B2M.* Data are representative of n=3 in (A) and n=4 in (B-E). Data (mean ± SEM) in panels A-E and were analysed using a One-way ANOVA with Tukey’s post hoc test for multiple comparisons. *P<0.05, **P<0.01, ***P<0.001 compared to *ATP13A3*^LK726X^.

**Supplement Figure 16.**
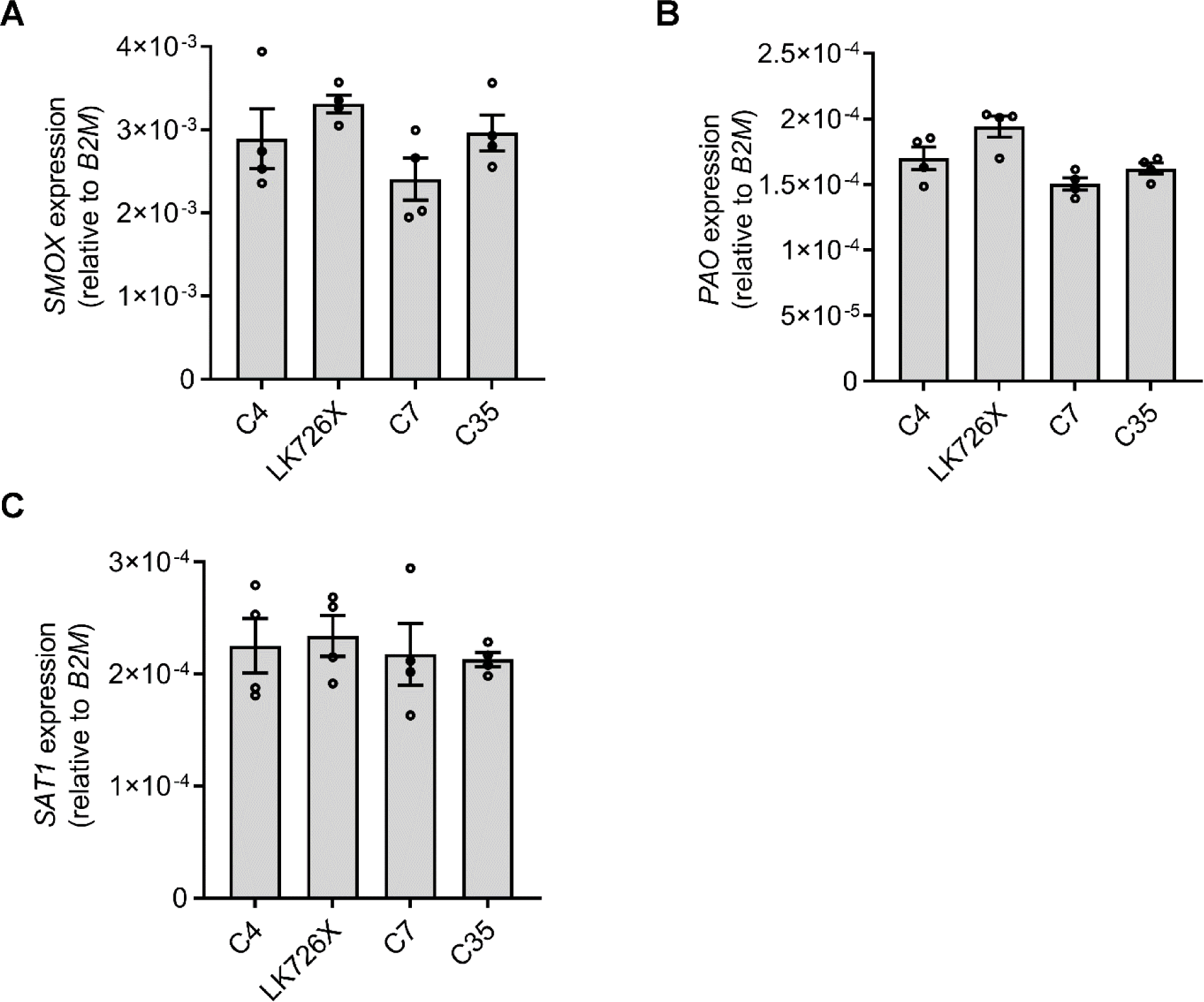
mRNA expression of polyamine catabolic enzymes in control and ATP13A3^LK726X^ BOECs. mRNA expression of (A) *SMOX* (B) *PAO* (C) *SAT1* in control BOECs (C4, C7, C35) and *ATP13A3*^LK726X^ (LK726X) BOECs. Data (n=4 experiments) are mean ± SEM and presented as expression relative to *B2M*.

**Supplement Figure 17.**
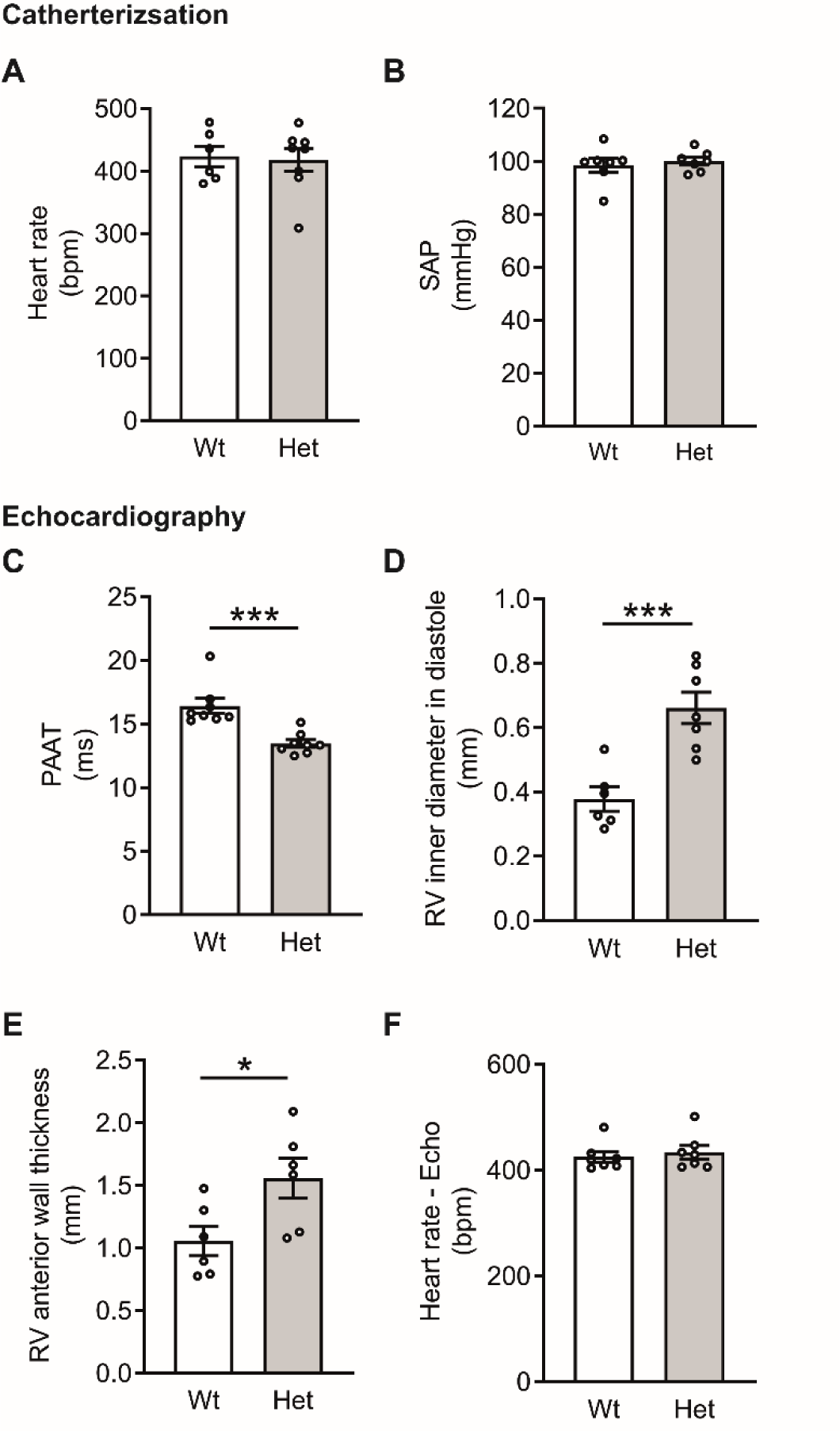
Mice harbouring an *Atp13a3*^P452Lfs^ mutation spontaneously develop PAH and right ventricular dysfunction. (A-B) Invasive hemodynamic measurement of (A) heart rate and (B) systolic blood pressure (SAP) *Atp13a3*^P452Lfs^ heterozygous mice and controls (n = 6, 10). (C-F) Ultrasound echocardiography assessment of cardiopulmonary function. (C) Pulmonary artery acceleration time, (D) Right ventricular inner diameter, (E) Right ventricular end diastolic anterior wall thickness and (F) heart rate. Data are mean ± SEM analysed using an unpaired two-tailed t-test with Welch’s correction. *P<0.05, ***P<0.001

**Supplement table 1.**
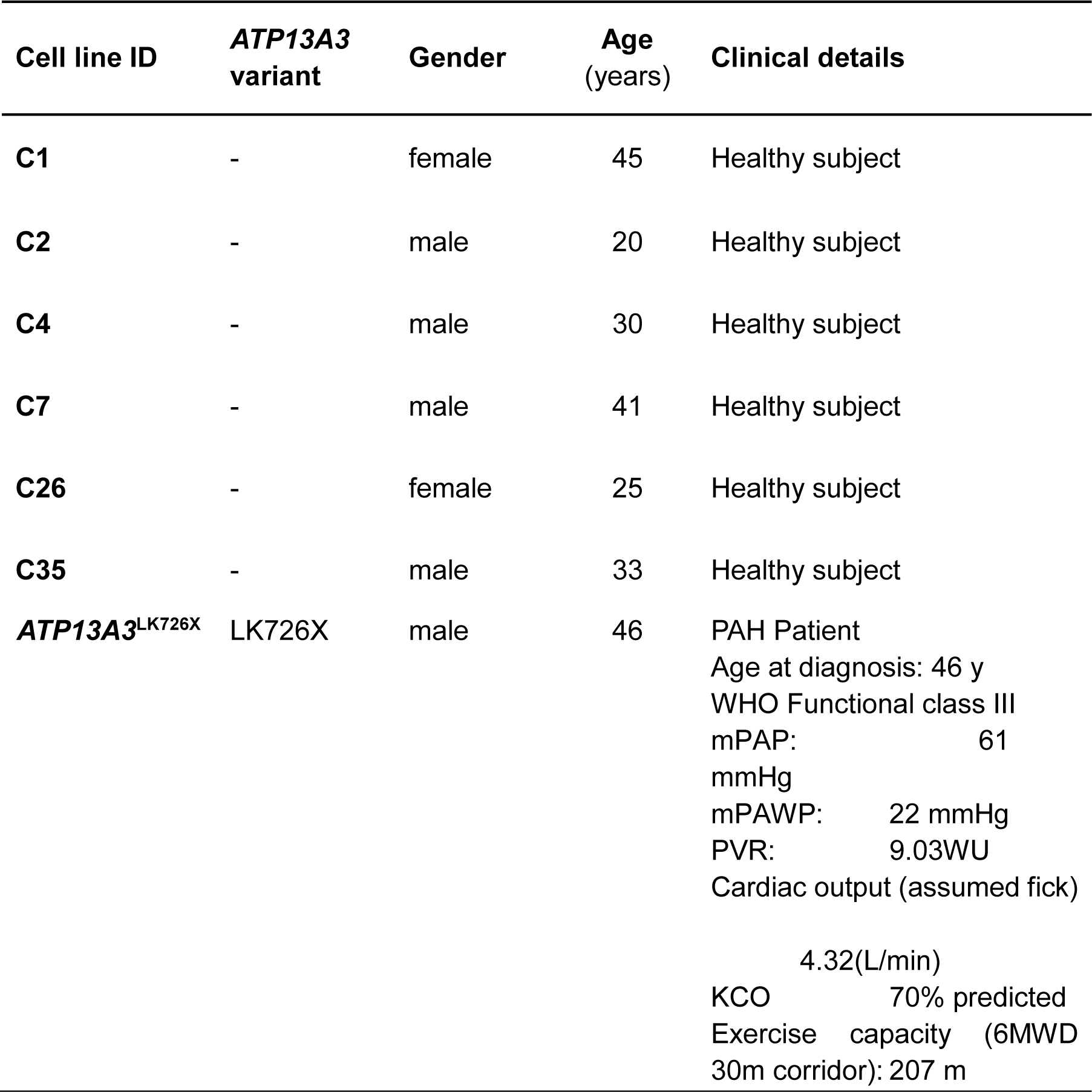
*ATP13A3* variant and demographic information of different BOEC lines.

| Cell line ID | *ATP13A3* variant | Gender | Age (years) | Clinical details |
| --- | --- | --- | --- | --- |
| **C1** | - | female | 45 | Healthy subject |
| **C2** | - | male | 20 | Healthy subject |
| **C4** | - | male | 30 | Healthy subject |
| **C7** | - | male | 41 | Healthy subject |
| **C26** | - | female | 25 | Healthy subject |
| **C35** | - | male | 33 | Healthy subject |
| ***ATP13A3*<sup>LK726X</sup>** | LK726X | male | 46 | PAH Patient<br>Age at diagnosis: 46 y<br>WHO Functional class III<br>mPAP: 61 mmHg<br>mPAWP: 22 mmHg<br>PVR: 9.03WU<br>Cardiac output (assumed fick)<br>4.32(L/min)<br>KCO 70% predicted<br>Exercise capacity (6MWD 30m corridor): 207 m |

**Supplement Table 2.**
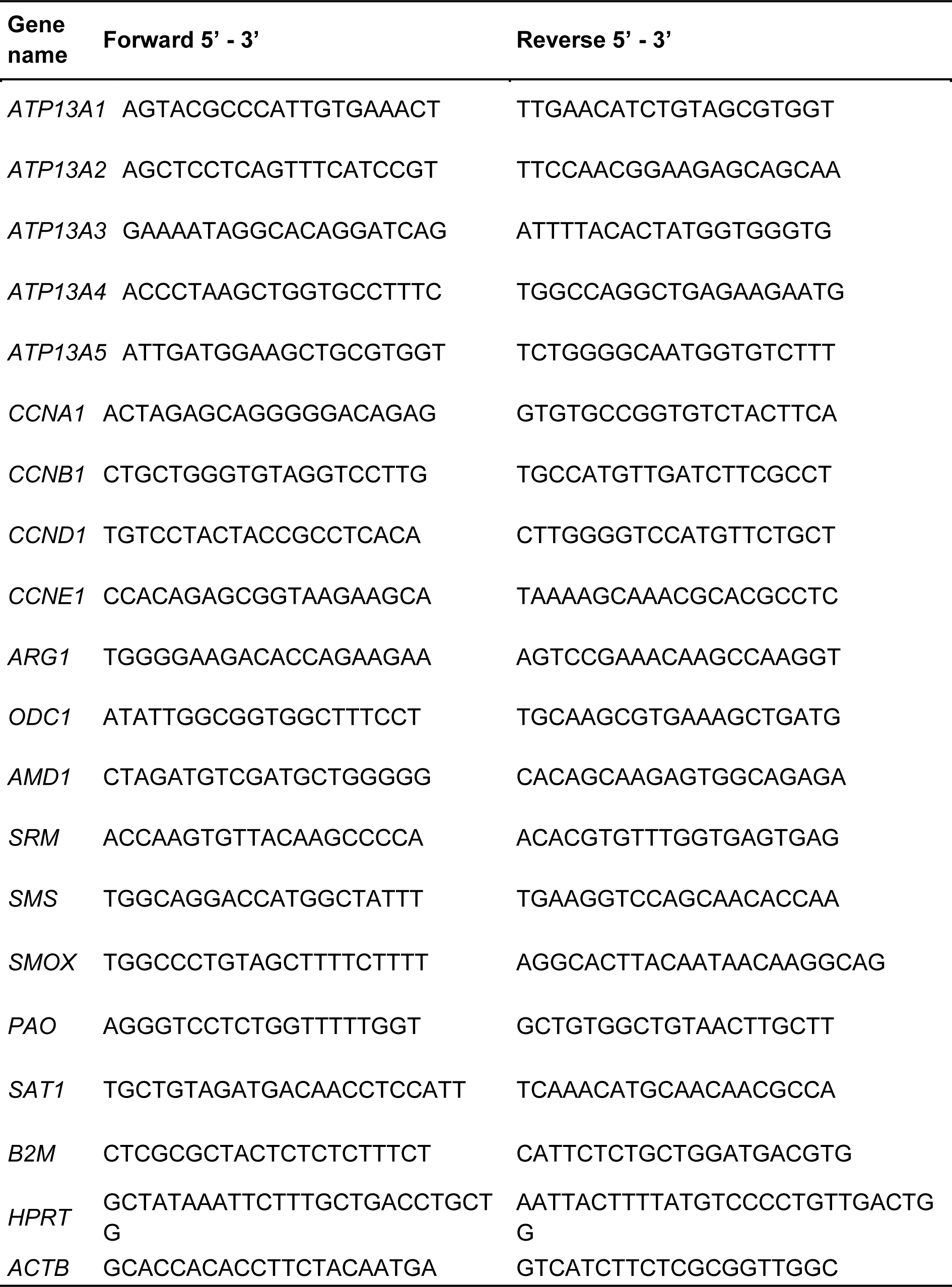
Primer sequences for human genes analysed using qPCR.

**Supplement Table 3.** antibodies used for cellular immunostaining.

| Antibody | Source | Dilution | Secondary | Dilution |
| --- | --- | --- | --- | --- |
| Rabbit anti-ATP13A3 | Sigma-Aldrich<br>HPA029471 | 1:200 | rabbit Alexa<br>Fluor®488 | 1:200 |
| Mouse Anti-VE-Cadherin CD144 | BD<br>Biosciences<br>555661 | 1:200 | Mouse<br>Alexa<br>Fluor®568 | 1:200 |
| Mouse Anti-EEA1 | BD<br>Biosciences<br>610456 | 1:200 | Mouse<br>Alexa<br>Fluor®568 | 1:200 |
| Mouse Anti-Rab7 | SANTA CRUZ<br>sc-376362 | 1:200 | Mouse<br>Alexa<br>Fluor®568 | 1:200 |
| Mouse Anti-Rab11a | SANTA CRUZ<br>Sc-166912 | 1:200 | Mouse<br>Alexa<br>Fluor®568 | 1:200 |
| Mouse Anti-LAMP1 | SANTA CRUZ<br>Sc-20011 | 1:200 | Mouse<br>Alexa<br>Fluor®568 | 1:200 |

